# The impact of artifact removal approaches on TMS–EEG signal

**DOI:** 10.1101/2021.01.15.426817

**Authors:** Giacomo Bertazzoli, Romina Esposito, Tuomas P. Mutanen, Clarissa Ferrari, Risto J. Ilmoniemi, Carlo Miniussi, Marta Bortoletto

## Abstract

Transcranial magnetic stimulation (TMS)-evoked potentials (TEPs) allow one to assess cortical excitability and effective connectivity in clinical and basic research. However, obtaining clean TEPs is challenging due to the various TMS-related artifacts that contaminate the electroencephalographic (EEG) signal when the TMS pulse is delivered. Different preprocessing approaches have been employed to remove the artifacts, but the degree of artifact reduction or signal distortion introduced in this phase of analysis is still unknown. Knowing and controlling this potential source of uncertainty will increase the inter-rater reliability of TEPs and improve the comparability between TMS–EEG studies. The goal of this study was to assess the variability in TEP waveforms due to of the use of different preprocessing pipelines. To accomplish this aim, we preprocessed the same TMS–EEG data with four different pipelines and compared the results. The dataset was obtained from 16 subjects in two identical recording sessions, each session consisting of both left dorsolateral prefrontal cortex and left inferior parietal lobule stimulation at 100% of the resting motor threshold. Considerable differences in TEP amplitudes were found between the preprocessing pipelines. Topographies of TEPs from the different pipelines were all highly correlated (ρ>0.8) at latencies over 100 ms. By contrast, waveforms at latencies under 100 ms showed a variable level of correlation, with ρ ranging between 0.2 and 0.9. Moreover, the test–retest reliability of TEPs depended on the preprocessing pipeline. Taken together, these results take us to suggest that the choice of the preprocessing approach has a marked impact on the final TEP, and that caution should be taken when comparing TMS–EEG studies that used different approaches. Finally, we propose strategies to control this source of variability.

## 1. Introduction

Over the last decade, transcranial magnetic stimulation (TMS) coupled with electroencephalography (EEG) has been an increasingly popular tool for investigating cortical excitability and connectivity. TMS-evoked potentials (TEPs), i.e., the cortical responses time-locked to the TMS pulse, provide information on the cortical excitability (Ilmoniemi and Kičić, 2010; Komssi and Kähkönen, 2006; Veniero et al., 2013) and effective connectivity, i.e., patterns of signal spread through the network in which the stimulated area is embedded (Bortoletto et al., 2015). TEPs can provide direct causal information about the status of the stimulated cortex and connected areas, without the similar need for *a priori* assumptions as with functional magnetic resonance imaging (fMRI) or EEG alone. For these reasons, TEPs are used in clinical research to investigate neurophysiological alterations related to several psychiatric and neurological diseases (for review, see Tremblay et al., 2019), e.g., in disorders of consciousness to distinguish between different clinical subtypes of patients (Rosanova et al., 2012) and in Alzheimer’s disease as a marker of disease progression (Bagattini et al., 2019). Therefore, TEPs have been proposed as biomarkers that could improve diagnosis and monitor treatment-induced neuronal changes.

A crucial step for the development of a useful biomarker is demonstrating high reliability, usually equated with reproducibility, i.e., the degree to which a measurement is free from variable measurement error (Mokkink et al., 2010). High reliability allows one to reliably detect meaningful changes in the signals of interest. Reliability can be assessed in a population under stable conditions by taking the same measure with different instruments (internal consistency), by different raters (inter-rater reliability) and by the same rater over time (intra-rater reliability or test–retest reliability) (Beaulieu et al., 2017; Mokkink et al., 2010).

Up to now, only test–retest reliability of TEPs has been assessed in a few studies, which found overall high reliability and replicability for TEP components (Casarotto et al., 2010; Lioumis et al., 2009), mainly at late latencies (Kerwin et al., 2018; Ozdemir et al., 2020).

However, these studies quantified the test–retest reliability using the same data-analysis pipeline, while reproducibility of TEPs may be affected by the preprocessing steps. Indeed, TMS–EEG signals are often heavily contaminated by various TMS-related artifacts, which can be an order of magnitude higher than the neuronal signal and time-locked with the TMS pulse, reducing the signal-to-noise ratio (SNR). To deal with this issue, several methodologies and algorithms have been developed to remove the artifacts while sparing the neuronal signal. On one side, the preprocessing phase allows the extraction of TEPs, but on the other side it could also affect their reproducibility. Specifically, the use of different approaches to reduce artefacts by different experimenters could affect the inter-rater reliability of TEPs. Moreover, the results of the test–retest reliability could vary across studies, due to the use of different approaches.

The goal of this study is to determine the amount of variability brought in the TEPs by the use of different preprocessing pipelines. To this end, we processed the same TMS–EEG dataset with four published pipelines: ARTIST (Wu et al., 2018), TMSEEG (Atluri et al., 2016), TESA (Rogasch et al., 2017), and SOUND/SSP–SIR (Mutanen et al., 2016, 2018), and compared the resulting TEPs. Although these pipelines share the common goal of removing artifacts while sparing the neuronal signal from TMS–EEG recordings, they employ different strategies. ARTIST is a fully automatic pipeline, developed to minimize the variability brought by the experimenter. Conversely, TMSEEG is operated via a fully manual graphical user interface (GUI) that guides the user through the steps needed to clean TMS–EEG signals. TESA, TMSEEG and ARTIST use two steps of independent component analysis (ICA) as a core function to isolate and remove artifacts, although employing different algorithms. SOUND-SSP–SIR employs ICA only for the removal of ocular artifacts, while its core functions are the source-estimate-utilizing noise-discarding algorithm (SOUND) (Mutanen et al., 2018) and signal-space projection–source-informed reconstruction (SSP–SIR) (Mutanen et al., 2016). Taken together, these processing differences may impact amplitude and topography of TEP components and ultimately their reproducibility.

## 2. Materials and methods

The dataset used in this study was acquired as a part of a larger TMS–EEG study to investigate brain connectivity. The study followed the Helsinki Declaration guidelines and was approved by the Human Research Ethical Committee of the University of Trento (protocol number 2017-014). Data from this study are available on [OpenNeuro]. Scripts regarding the preprocessing pipelines from this study are available on [GitHub].

### 2.1 Participants

Sixteen healthy volunteers (age 24.5 ± 2.8, 7 females) were enrolled in the experiment. Each participant gave a written informed consent for participating in this study and was screened for MRI and TMS compatibility (Rossi et al., 2009; Sammet, 2016). Each subject underwent three experimental sessions: in the first session, MRI and fMRI were acquired to obtain individualized structural and functional localization of the default-mode network (DMN); the other two identical sessions (test–retest) were 72.3 ± 35.8 days apart and involved TMS–EEG coregistration during DMN stimulation.

### 2.2 MRI data acquisition

High-resolution anatomical images were acquired with two *T*1-weighted anatomical scans (MP-RAGE; 1×1×1 mm^3^; FOV, 256224 mm^2^; 176 slices; GRAPPA acquisition with an acceleration factor of 2; TR, 2700/ 2500 ms; TE, 4.18/ 3.37 ms; inversion time (TI), 1020/ 1200 ms; 7° / 12° flip angle), through a 4T Siemens MedSpec Synco MR scanner and a birdcage transmit 8-channel receive head radiofrequency coil. Single shot *T*2*-weighted gradient-recalled echo-planar imaging (EPI) sequence was used to acquire functional images. A 30-slice protocol was used, acquiring each slice in ascending interleaved order, within a repletion time (TR) of 2000 ms (voxel resolution, 3×3×3 mm^3^; echo time (TE), 28 ms; flip angle (FA), 73°; field of view (FOV), 192×192 mm^2^). Each run consisted of 200 volumes.

### 2.3 TMS protocol and targeting

Single-pulse TMS was delivered through a Magstim Rapid magnetic stimulator (Magstim, Whitland, UK) and a 70-mm figure-of-eight coil. In each session, the coil location corresponding to the hotspot for the right first dorsal interosseous muscle stimulation was identified and the resting motor threshold (rMT) was measured as the lowest intensity producing motor-evoked potentials of over 50µV in a minimum of 5 out of 10 trials in the electromyogram recorded with bipolar montage (Rossini et al., 2015). Thereafter, the stimulator intensity was set at 100% of the rMT. Four nodes of the DMN extracted from functional maps (see supplementary materials for details) were stimulated: left and right dorsolateral prefrontal cortex (DLPFC) and left and right inferior parietal lobule (IPL). The Talairach coordinates of the point of local maxima in each node and the structural MRI images were input to the SofTaxic Neuronavigation system (E.M.S., Bologna, Italy) to guide an accurate and consistent positioning of the TMS coil. The order of stimulation target was randomized across participants. On each site, 120 biphasic TMS pulses were delivered with an inter-stimulus interval of 2–10 seconds. The coil was placed by the experimenter in a position tangential to the scalp, with an angle of approximately 45° between the interhemispheric sulcus and the coil handle, with the handle pointing backwards, for the DLPFC and 10° for the IPL.

For the purposes of this study, we limited our analyses to the left DLPFC and the left IPL, from both the test (T1) and retest session (T2).

### 2.4 EEG data acquisition

EEG signals were acquired with a BrainAmp DC TMS-compatible system (BrainProducts GmbH, Germany), with a continuous recording from BrainCap 62 TMS-compatible Ag/AgCl-coated Multitrode electrodes (BrainProducts GmbH, Germany) positioned on the scalp according to the 10/10 International System. One electrode was placed to the left-eye temporal canthus to detect horizontal eye movements, and another electrode was placed beneath the left eye to detect vertical eye movements and blinks. TP9 was used as online reference. The ground electrode was placed at FPz. Signals from all channels were band-pass filtered at 0.1–1000 Hz and digitized at the sampling rate of 5 kHz. The skin–electrode impedance was kept below 5 kΩ. Trigger marks at the TMS pulse delivery were sent to the EEG system using a customized MATLAB script (MathWorks^®^, Inc., Massachusetts, USA).

### 2.5 EEG signal preprocessing

#### Preprocessing pipelines

Next, we give a brief introduction to the preprocessing pipelines employed in this study. For a detailed description, please refer to the respective original papers, i.e., ARTIST (Wu et al., 2018), TMSEEG (Atluri et al., 2016), TESA (Rogasch et al., 2017), SOUND/SSP–SIR (Mutanen et al., 2018, 2016).

#### 2.5.1 TMSEEG

The TMSEEG (Atluri et al., 2016) pipeline is run via a GUI from which the user can call, in a fixed order, all the functions needed to process the TMS–EEG signal. It was designed to provide an easy-to-use streamline to guide the user (novice or advanced) through all the steps needed to clean raw TMS–EEG data. TMSEEG is an ICA-based pipeline (*fastICA* algorithm, Hyvärinen and Oja, 2000), with two ICA steps: the first aims at removing the large-amplitude artifacts, while the second removes the other TMS-related and common EEG artifacts. TMSEEG is fully manual: trials/channels and ICA-components’ rejection are choices made by the user.

#### 2.5.2 *TMS–EEG signal analyser*, TESA

The TESA (Rogasch et al., 2017) toolbox is a set of functions that works as an extension of the EEGlab software (Delorme and Makeig, 2004). Here the user can choose to run the functions from the EEG lab GUI or directly script the pipeline, calling the functions from MATLAB command window. The TESA toolbox was designed to provide multiple tools to the user that could be combined and used as preferred. For this reason, it has multiple core functions to remove artifacts from the signal (ICA, principal component analysis, enhanced deflation method, etc.); therefore, it does not have a unique pipeline. In this work, we chose the default pipeline suggested in (Rogasch et al., 2017). In this case, TESA is used to build an ICA-based pipeline (*fastICA* algorithm), with two ICA steps: the first aims to remove the large-amplitude artifacts (TMS-related muscle, decay and movement) and the second aims to remove the remaining artifacts. TESA can be used manually through the EEGlab GUI, semi-automatically (using automatic bad trials/channels rejection and supervised ICA-component rejection) or automatically (automatic bad trials/channels and ICA-component rejection), calling the functions through MATLAB command window. We employed the semi-automatic mode.

#### 2.5.3 *Automated aRTIfact rejection for Single-pulse TMS–EEG Data*, ARTIST

The idea behind ARTIST (Wu et al., 2018) is to remove the variability introduced by the user, by delegating most of the choices to an automatic algorithm. The user sets the necessary parameters for the cleaning (epoch length, TMS-pulse interpolation period etc.) and calls the main function from the MATLAB command window. The algorithm then performs all the necessary steps automatically. Like TMSEEG and TESA, it is an ICA-based pipeline (*Infomax* algorithm (Makeig et al., 1997)). It employs the first ICA run to remove the big decay artifact and the second ICA run to remove the remaining artifactual data. Bad trials/channels and ICA-components are labelled and rejected by specific algorithms and trained classifiers.

#### 2.5.4 *Source-estimate-utilizing noise-discarding algorithm* (SOUND) *and signal-space projection–source-informed reconstruction* (SSP–SIR)

The SOUND-SSP–SIR approach substitutes ICA largely with the SOUND (Mutanen et al., 2018) and SSP–SIR (Mutanen et al., 2016) algorithms as core functions to remove TMS–EEG artifacts. The SOUND algorithm reconstructs the signal in the source volume. Then, it attenuates artifactual segments of data in each sensor, by estimating how well that segment of signal is predicted by all other sensors. This process runs automatically and does not require the user to choose the signal to be removed. SSP–SIR, instead, is specifically designed to attenuate the TMS-related muscle artifacts, using their time-frequency features. The signal components that capture the TMS-related muscle activity can be rejected manually by the user, or automatically by setting an appropriate threshold. We chose to use SSP–SIR manually. The SOUND-SSP–SIR approach, applied here, is a combination of methods that have been unified in a coherent pipeline in previous studies (Bagattini et al., 2019; Bortoletto et al., 2020). Finally, an ICA step is also present in this pipeline (*Infomax* algorithm), but it is confined to removing ocular artifacts as these can be assumed to appear relatively independently from TMS-evoked brain signals. The SOUND-SSP–SIR pipeline does not have a GUI and it can be used by calling the functions through the MATLAB command window.

##### 2.5.5 Preprocessing parameters and steps

Raw TMS–EEG signals were processed with the above-mentioned pipelines. Table 1 illustrates the workflow for each pipeline. Epoching was set to -/+1 second. Electrooculography (EOG) electrodes were removed using the built-in EEGlab function for channel removal. Given that TMS-pulse artifact lasts up to 5 ms with the employed recording settings (Veniero et al., 2009), data were removed and interpolated from –1 to 6 ms. Data were downsampled to 1000 Hz. When suggested, demeaning was applied using the whole epoch (TMSEEG and TESA). Baseline correction was set from –1000 to –2 ms. Channel and trial rejection was done either manually (TMSEEG) or automatically (ARTIST and TESA) employing built-in functions in EEGlab (Delorme and Makeig, 2004). The SOUND-SSP–SIR pipeline requires only manual trial rejection, as bad channels are automatically corrected by the SOUND step. Data were bandpass filtered at 1–90 Hz and notch filtered at 48–52 Hz. When high-pass filtering was applied alone, the passband started at 1 Hz. Rereferencing was applied using the average of all electrodes. ICA and SSP–SIR component selection was performed either manually (TMSEEG, TESA, SOUND-SSP–SIR) or automatically (ARTIST). When run manually, ICA component selection was done independently by authors G.B. and R.E. and SSP–SIR component selection by G.B. and M.B. Components that were selected by just one author were discussed and a joint decision was made. For statistical analysis, the epoch was reduced to –100…+350 ms and baseline correction from –100 to –2 ms was performed.

**Table 1:** Steps order in each processing pipeline. Exact values for each parameter are described in the “Preprocessing parameters and steps” paragraph.

|  | **ARTIST** | **TMSEEG** | **TESA** | **SOUND-SSP-SIR** |
| --- | --- | --- | --- | --- |
| **1** | EOGs removal | Epoching | EOGs removal | EOGs removal |
| **2** | TMS-pulse interpolation | Demeaning <sup>#</sup> | Channel rejection <sup>+</sup> | High-pass filter |
| **3** | Downsampling) | EOGs removal | Epoching | Epoching |
| **4** | Detrending | TMS-pulse removal* | Demeaning <sup>#</sup> | TMS-pulse interpolation |
| **5** | Epoching | Channels and trials rejection -1 | TMS-pulse interpolation | Baseline correction |
| **6** | **ICA -1** | **ICA -1** | Downsampling | **SOUND** |
| **7** | Bandpass – Notch filters | Bandpass – Notch filters | Trials rejection <sup>+</sup> | Trials rejection |
| **8** | Channels and trials rejection <sup>+</sup> | **ICA -2** | **ICA -1** <sup>§</sup> | ICA <sup>@</sup> |
| **9** | Bad-channels interpolation | Channels and trials rejection - 2 | Bandpass – Notch filters | **SSP-SIR** <sup>§</sup> |
| **10** | **ICA -2** | Rereferencing | **ICA -2** <sup>§</sup> | Bandpass – Notch filters |
| 11 | Rereferencing | Bad-channels interpolation | Bad-channels interpolation | Rereferencing |
| 12 | Baseline correction | TMS-pulse interpolation* | Rereferencing | Downsampling |
| 13 |  | Baseline correction <sup>×</sup> | Baseline correction | Baseline correction |
| 14 |  | Downsampling <sup>×</sup> |  |  |

\* TMS-pulse artifact was removed, but not interpolated. Instead, the signal was cut and pasted. In the end of the pipeline, the removed TMS-pulse latencies were replaced with NaNs and then interpolated.

### 2.6 Statistical analysis

First, we investigated the between-pipelines differences in trials, channels and ICA components removed during the preprocessing. Channels rejected and components removed in the ICA were summarized together using the data rank. The data rank represents the number of independent components that can be individuated in the data. The higher the rank, the “richer” are the data in terms of sources that compose the signal. In raw TMS–EEG signal, the rank is equal to the number of recording channels and it decreases by one unit during the preprocessing for each channel interpolated and ICA-component removed (minus one for the rereferencing). Depending on the preprocessing strategy, different pipelines interpolate a different number of bad channels and remove or attenuate a different number of components, hence impacting the rank in different ways. With methods like ICA or SSP, the decrease in rank corresponds to exactly the number of removed artifact components (signal dimensions). SOUND, however, weighs different signal dimensions based on the noise estimates. This means that while SOUND theoretically conserves the rank, it can effectively squash the noisiest dimensions close to zero. To allow a fair comparison between the different pipelines, we estimated the effective rank of the data as follows: We formed the singular-value decomposition of the data and defined the effective rank as the number of dimensions having singular value greater than 1% of the biggest singular value. Differences in data rank after the preprocessing were assessed with a separated one-way ANOVA per each condition (IPL session 1, IPL session 2, DLPFC session 1 and DLPFC session 2). Similarly, differences in trial removal were assessed with a Kruskal–Wallis test, to account for the non-normal distribution of the data. Post-hoc tests were applied when appropriate with alpha threshold 0.0042, for a series of six two-tailed tests with Bonferroni correction.

Reproducibility, i.e., the reliability of TEPs derived by different preprocessing pipelines, was assessed by testing for (1) differences of the final TEPs, (2) similarities of the final TEPs, and (3) comparing the test–retest agreement between TEP components in session 1 and session 2 across pipelines.

Differences across TEPs were investigated separately for the two areas (IPL, DLPFC) and sessions 1–2; in total, four separate one-way repeated measure ANOVAs with “preprocessing pipeline” as a four-level factor were used to test the hypothesis that there were differences in TEP amplitudes across preprocessing pipelines in some channels and time points after the TMS pulse. For controlling the familywise error rate, we used cluster-based permutation tests (Oostenveld et al., 2011). This strategy is commonly used when there is no a priori hypothesis on where or when the effect will be. The ANOVA with cluster-based correction was ran at each time point after the TMS-pulse interpolation (6 to 350 ms, SR: 1000 Hz) in each channel (63). Time points with significant test statistics (alpha level was set at 0.05) were clustered together, constrained to those that involved at least 3 neighboring channels. Post-hoc tests were applied as a series of cluster-based two-tailed *t*-tests for paired samples between all possible combinations of pipelines, with alpha < 0.0042 (0.025 divided for six comparisons) to correct for multiple comparisons.

Similarities across the TEPs, produced by different pipelines, were investigated through pairwise Spearman correlations in both temporal and spatial dimensions. The spatial correlation of TEPs for each pair of preprocessing pipelines was computed at each time point, by correlating voltage values of all channels in a specific pair of conditions, i.e., two preprocessing pipelines, for each individual. The resulting correlation values were then *z*-transformed and averaged across subjects. The *z*-coefficients were tested at each time point against zero using a two-tailed *t*-test, to assess the significance of the correlation. For each test, alpha level was set at 0.025 (two-tailed *t*-test). Since each contrast comprised a series of 344 *t*-tests (one for each time point) we controlled the False Discovery Rate (FDR) with the Benjamini–Hochberg correction (Benjamini and Hochberg, 1995; Martínez-Cagigal, 2020). The average *z*-coefficients were transformed back into correlation values using the inverse Fisher *z*-transformation.

The temporal correlation of two TEPs was computed at each channel with the same logic of the spatial correlation, i.e., by correlating the voltage values in a specific interval between two conditions. Intervals were the whole epoch (–100 +350 ms) and smaller time windows (6–80 ms, 81–150 ms and 151–350 ms).

Lastly, we assessed how the test–retest reliability of the TEPs would change depending on the preprocessing pipeline used to clean the raw signal. We measured the test–retest reliability between sessions 1 and 2, for each preprocessing pipeline, through the concordance correlation coefficient (CCC). CCC measures the agreement between two variables *x* and *y*, i.e., the test and retest measures, as distance from the line *x*=*y* (Kerwin et al., 2018; King et al., 2007; Lin, 1989). Therefore, for this analysis, TEP components were individuated and their peak amplitudes and latencies were extracted following the collapsed localizer strategy suggested in Luck 2014 (Luck, 2014). CCC was computed between sessions 1 and 2 for both peak-amplitude and peak-latency values. CCCs were considered significantly different from zero when their bootstrapped 95% confidence intervals (CI) did not include zero. Differences across preprocessing pipelines’ CCCs were assessed using the percentile bootstrap method (Wilcox, 2009) within each peak. This method involves the calculation of the bootstrap distribution for the CCC difference between two conditions with its relative CI. If the CI of the CCC difference does not contain the zero, then the two conditions’ CCCs are significantly different.

## 3. Results

Figure 1 shows TEPs with relative topographies after being processed with the four pipelines, in both IPL (Fig. 1A) and DLPFC (Fig. 1B) for session 1. TEPs for session 2 are shown in supplementary results (Fig. 1S).

**Fig. 1:**
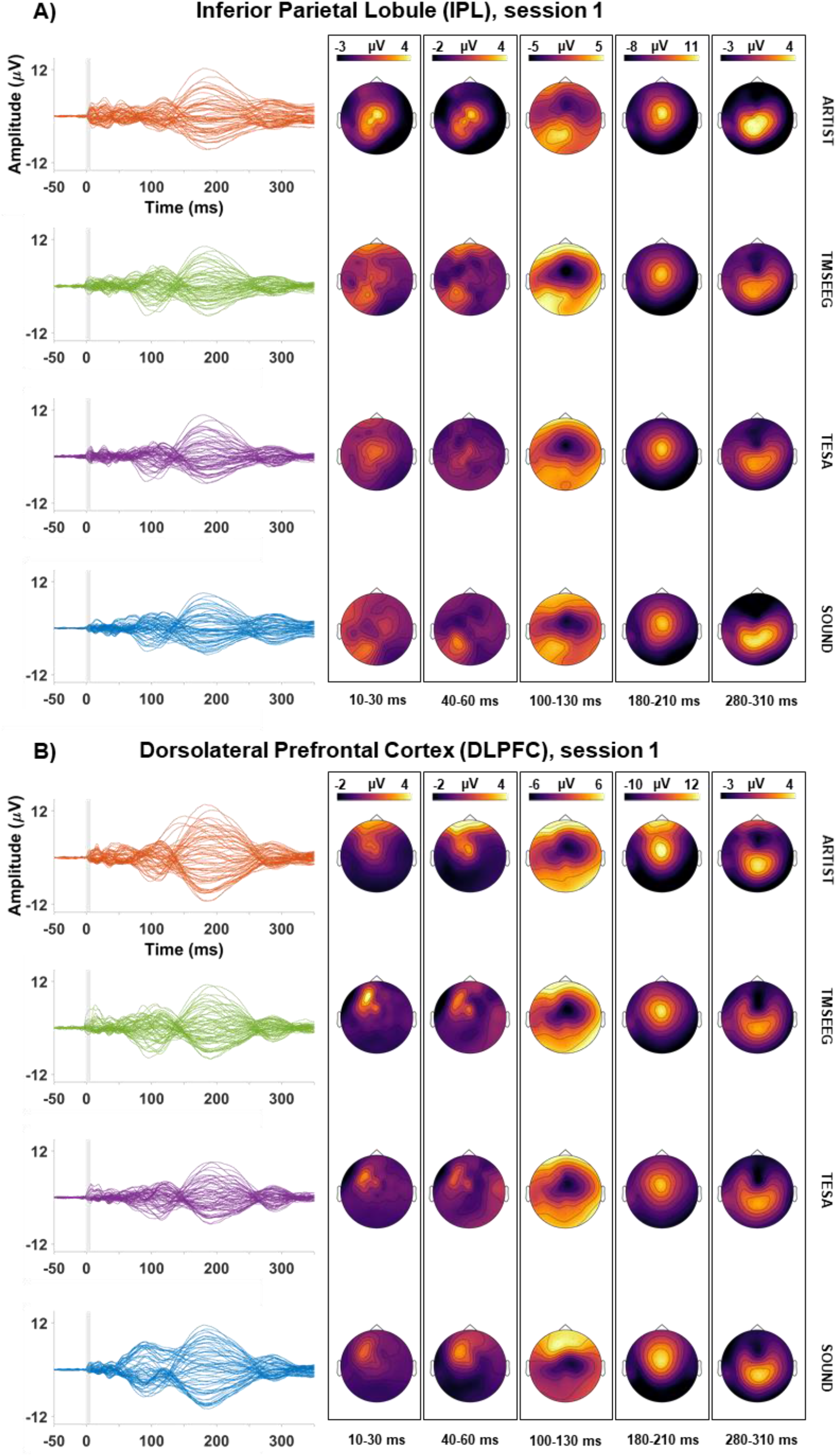
TEPs recorded after IPL (A) and DLPFC (B), processed with different pipelines (rows). The leftmost column depicts TEPs time-series, color-coded for each preprocessing pipeline. The other columns represent the scalp topographies of TEPs in five selected intervals. The topographies represent the mean voltage on the scalp in each interval. “SOUND” refers at the full SOUND-SSP–SIR pipeline.

The motor threshold did not significantly differ between the two sessions (session 1: 66.3 ± 8.8%, session 2: 66.4 ± 8.2% of the maximum stimulator output). The Kruskal–Wallis test on of the number of removed trials in session 1 (Table 1S) showed significant differences for IPL (χ^2^ = 13.7499, *p* = 0.0033) and DLPFC (χ^2^ = 12.7422, *p* = 0.0052). Post-hoc tests revealed significant differences (*p* threshold = 0.0042) for DLPFC between TMSEEG and SOUND-SSP–SIR (*t*= 23.2813, *p*= 0.0022). Data rank in session 1 was significantly different across pipelines for IPL (*F*=135.884, *p* < 0.001) and for DLPFC (*F*= 182.728, *p* < 0.001). Post-hoc analyses showed that the rank for ARTIST was significantly smaller than the rank for the other pipelines, while TMSEEG rank was significantly higher than the others (Fig. 2 and Table 2S). Trial and rank analysis for session 2 are shown in the supplementary materials (Fig. 2S and Table 2S).

**Table 2:** IPL post-hoc paired t-tests, session 1

| Comparison | Cluster Polarity | Cluster p | Cluster Latency (s) |
| --- | --- | --- | --- |
| ARTIST vs TMSEEG | + | < 0.001 | 0.007 to 0.350 |
|  | - | < 0.001 | 0.015 to 0.350 |
| ARTIST vs TESA | + | < 0.001 | 0.009 to 0.350 |
|  | - | 0.002 | 0.147 to 0.350 |
| ARTIST vs SOUND-SSP-SIR | + | < 0.001 | 0.006 to 0.350 |
|  | - | < 0.001 | 0.172 to 0.350 |
| TMSEEG vs TESA | + | 0.002 | 0.010 to 0.343 |
| TMSEEG vs SOUND-SSP-SIR | + | < 0.001 | 0.173 to 0.350 |
|  | - | < 0.001 | 0.191 to 0.350 |
| TESA vs SOUND-SSP-SIR | + | 0.004 | 0.224 to 0.350 |
|  | - | < 0.001 | 0.134 to 0.350 |

**Fig. 2:**
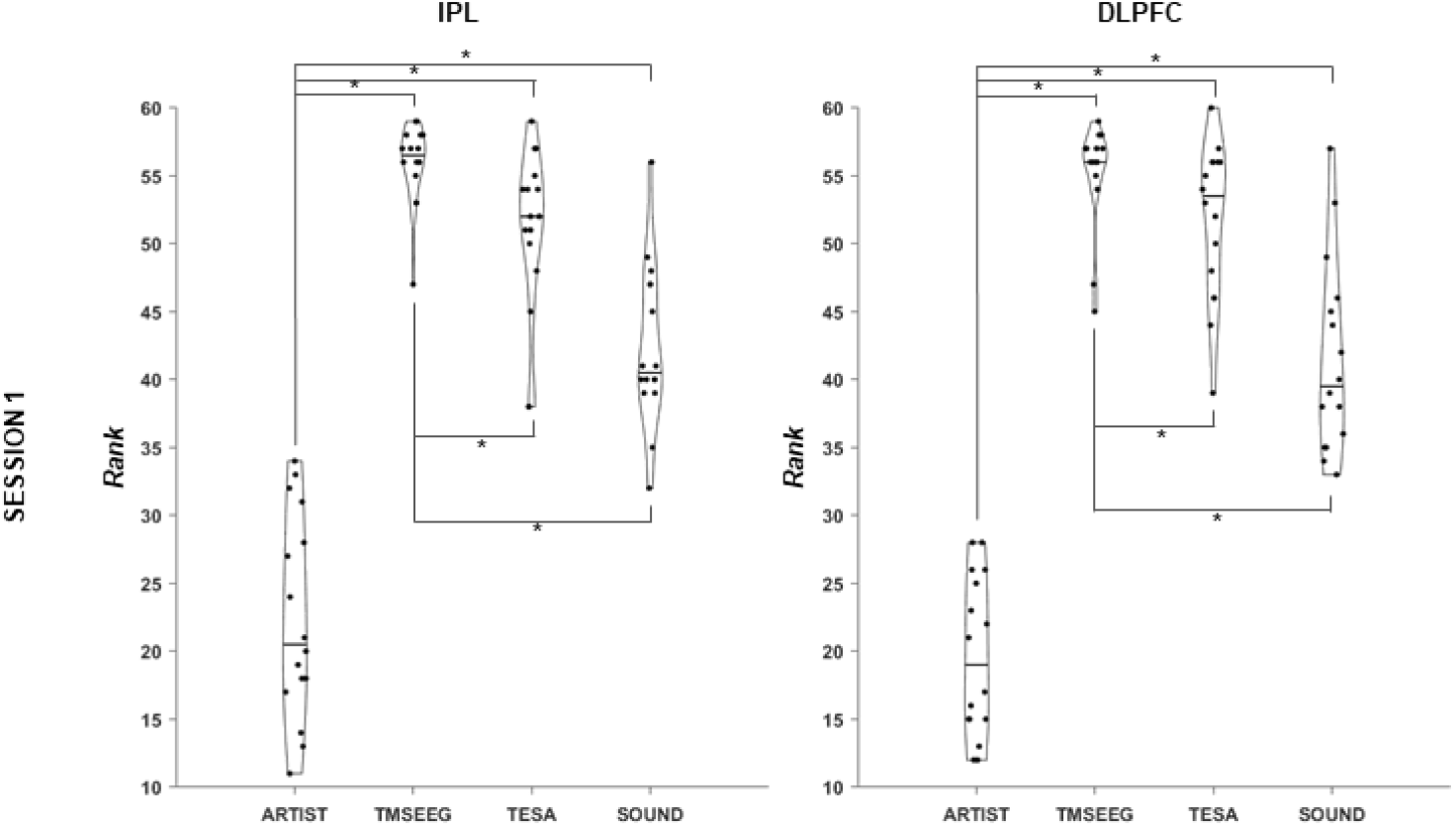
Data rank after each preprocessing pipeline for IPL and for DLPFC in session 1. Each violin plot depicts the distribution of individual subject rank after the preprocessing. Horizontal black lines represent the median. Black lines, with starts asterisks, connects conditions significantly different in the post-hoc analysis (p threshold= 0.0042). “SOUND” refers at the full SOUND-SSP–SIR pipeline.

### 3.1 Differences in TEPs amplitude

In IPL, TEPs derived from the four preprocessing pipelines were significantly different in amplitude (IPL: *p* =0.001, cluster window = 6–350 ms), as shown by the cluster-based ANOVA. In post-hoc tests (Fig. 3), differences were found in all contrasts and spanned for the whole epoch for all the comparisons with ARTIST and TMSEEG vs TESA. In TMSEEG vs SOUND-SSP–SIR and TESA vs SOUND-SSP–SIR the differences remained confined to late latencies (Table 2).

**Fig. 3:**
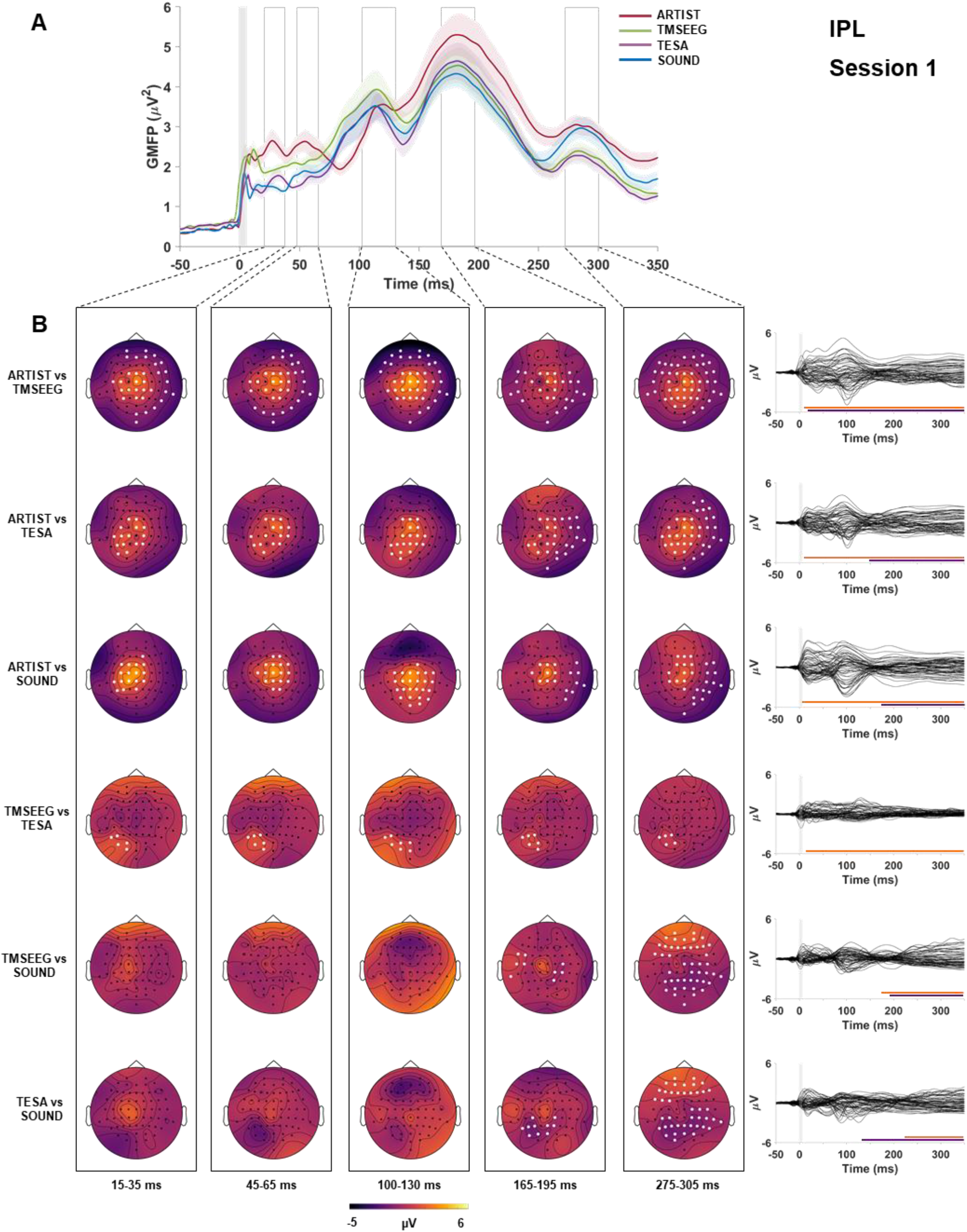
A: Global Mean Field Power (GMFP, y axes) over time (ms, x axis) of TEPs resulting from IPL stimulation, cleaned with the four preprocessing pipelines (color-coded). Shaded area around each colored line represents SEM. Shaded grey column around zero represents the TMS-pulse interpolation interval. B: scalp topographies of the voltage differences (color-coded) for each condition contrast (rows) in five selected time windows (columns). White dots represent significant channels. The rightmost column represents the time-series of the voltage differences over time in each contrast. Colored bars at the bottom of each time-series represent the temporal extend of significant cluster(s). Orange positive clusters, purple negative cluster. “SOUND” refers at the full SOUND-SSP–SIR pipeline.

Similarly, TEPs for DLPFC stimulation were different across processing pipelines (DLPFC: *p* =0.001, cluster window = 26–350 ms), as shown by the cluster-based ANOVA. Post-hoc tests (Fig. 4) showed significant differences confined at latencies above 100 ms, in four contrasts over six. The contrasts between TMSEEG vs TESA and TESA vs SOUND did not show significant differences (Table 3).

**Table 3:**
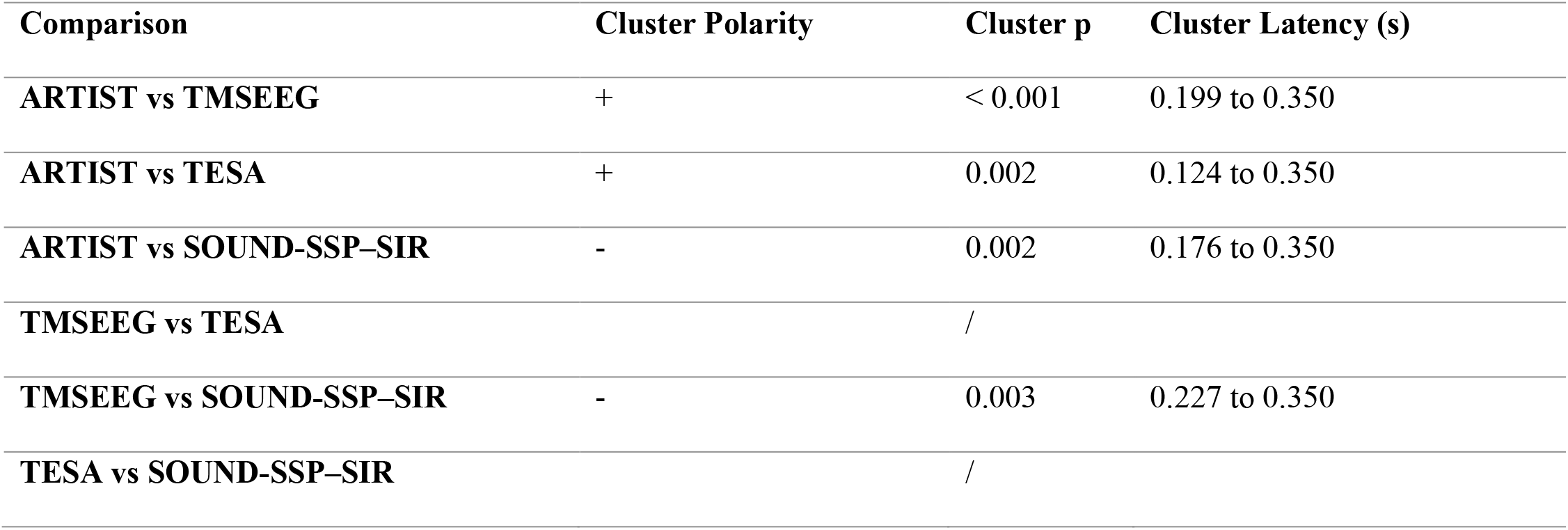
DLPFC post-hoc paired t-tests, session 1

| Comparison | Cluster Polarity | Cluster p | Cluster Latency (s) |
| --- | --- | --- | --- |
| ARTIST vs TMSEEG | + | < 0.001 | 0.199 to 0.350 |
| ARTIST vs TESA | + | 0.002 | 0.124 to 0.350 |
| ARTIST vs SOUND-SSP-SIR | - | 0.002 | 0.176 to 0.350 |
| TMSEEG vs TESA |  | / |  |
| TMSEEG vs SOUND-SSP-SIR | - | 0.003 | 0.227 to 0.350 |
| TESA vs SOUND-SSP-SIR |  | / |  |

**Fig. 4:**
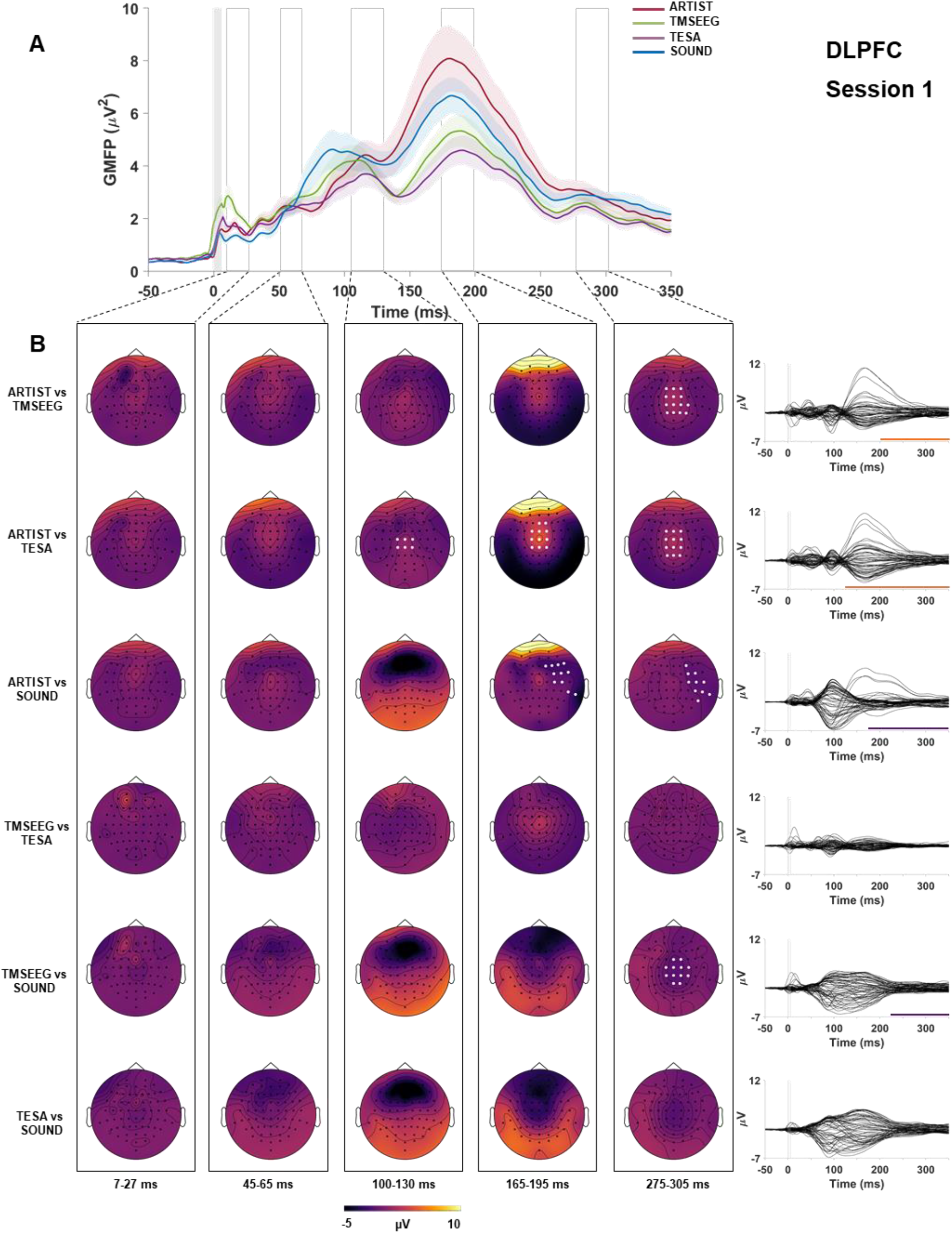
A: Global Mean Field Power (GMFP, y axes) over time (ms, x axis) of TEPs resulting from DLPFC stimulation, cleaned with the four preprocessing pipelines (color-coded). Shaded area around each colored line represents SEM. Shaded grey column around zero represents the TMS-pulse interpolation interval. B: scalp topographies of the voltage differences (color-coded) for each condition contrast (rows) in five selected time windows (columns). White dots represent significant channels (p threshold = 0.0042, cluster based corrected). The rightmost column represents the time-series of the voltage differences over time in each contrast. Colored bars at the bottom of each time-series represent the temporal extend of significant cluster(s). Orange positive clusters, purple negative cluster. “SOUND” refers at the full SOUND-SSP–SIR pipeline.

ANOVAs with related post-hocs between TEPs for session 2 are shown in the supplementary materials (Fig. 3S-4S, Tables 3S-4S).

### 3.2 TEP correlations

For interpretation of correlation values, we employed the scale of Shrout et al. 1998 (Shrout, 1998) (Table 5S).

Despite the marked differences in the cluster-based *t*-tests, TEPs across pipelines were positively correlated in most of the time points of the epoch in both IPL and DLPFC (Fig. 5-6 A). As expected, spatial correlations in the baseline period were high for both areas (moderate-to-substantial, ρ = 0.6–0.9), since all preprocessing pipelines shared the same raw signal. Spatial correlation dropped around the TMS pulse, and then slowly recovered. Correlation values latencies under 100 ms were the most variable across comparisons ranging from approximately ρ = 0.2 – 0.9, while correlations at latencies over 100 ms were consistently moderate-to-substantial (ρ>0.6) in all pairs.

**Fig. 5:**
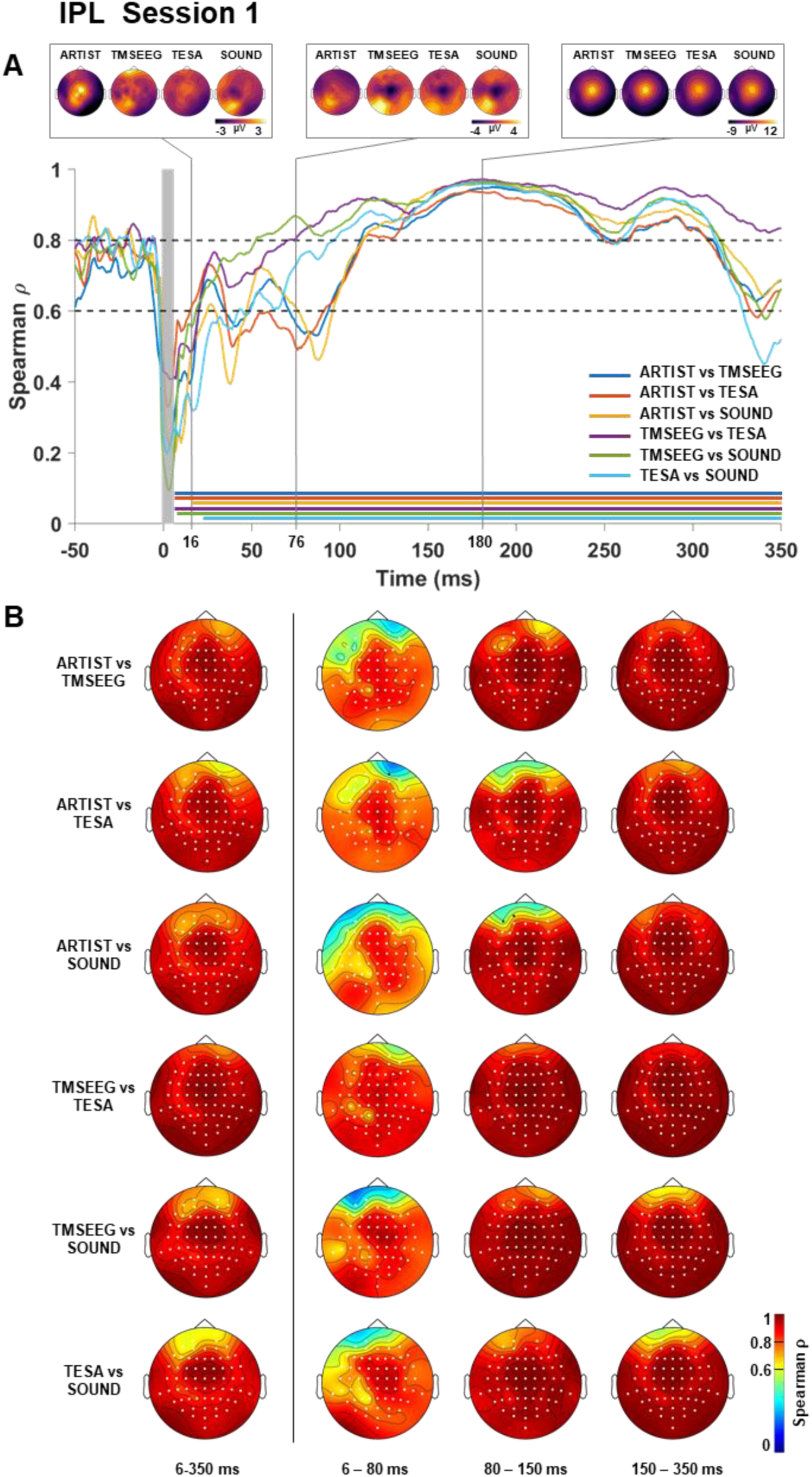
IPL, TEPs spatial and temporal correlation. (A) Each colored line represents the spatial correlation (y axis) over time (x axis) of all condition contrasts. Shaded grey column around zero represent the TMS-pulse interpolation interval. Horizontal dotted line represent threshold for moderate (0.6) and substantial (0.8) correlation, according to Shrout et al. 1998. Horizontal colored lines represent instant in time in which the correlation resulted significantly different from zero (p threshold = 0.0042, FDR corrected). On the top, are depicted three representative instantaneous scalp topographies in the four conditions. Voltage on the scalp topographies is color-coded. (B) Temporal correlation of each contrast (rows) in four time intervals (columns). Correlation values are color-coded. Channels significantly different from zero are highlighted in white (p threshold = 0.0042, FDR corrected). “SOUND” refers at the full SOUND-SSP–SIR pipeline.

**Fig. 6:**
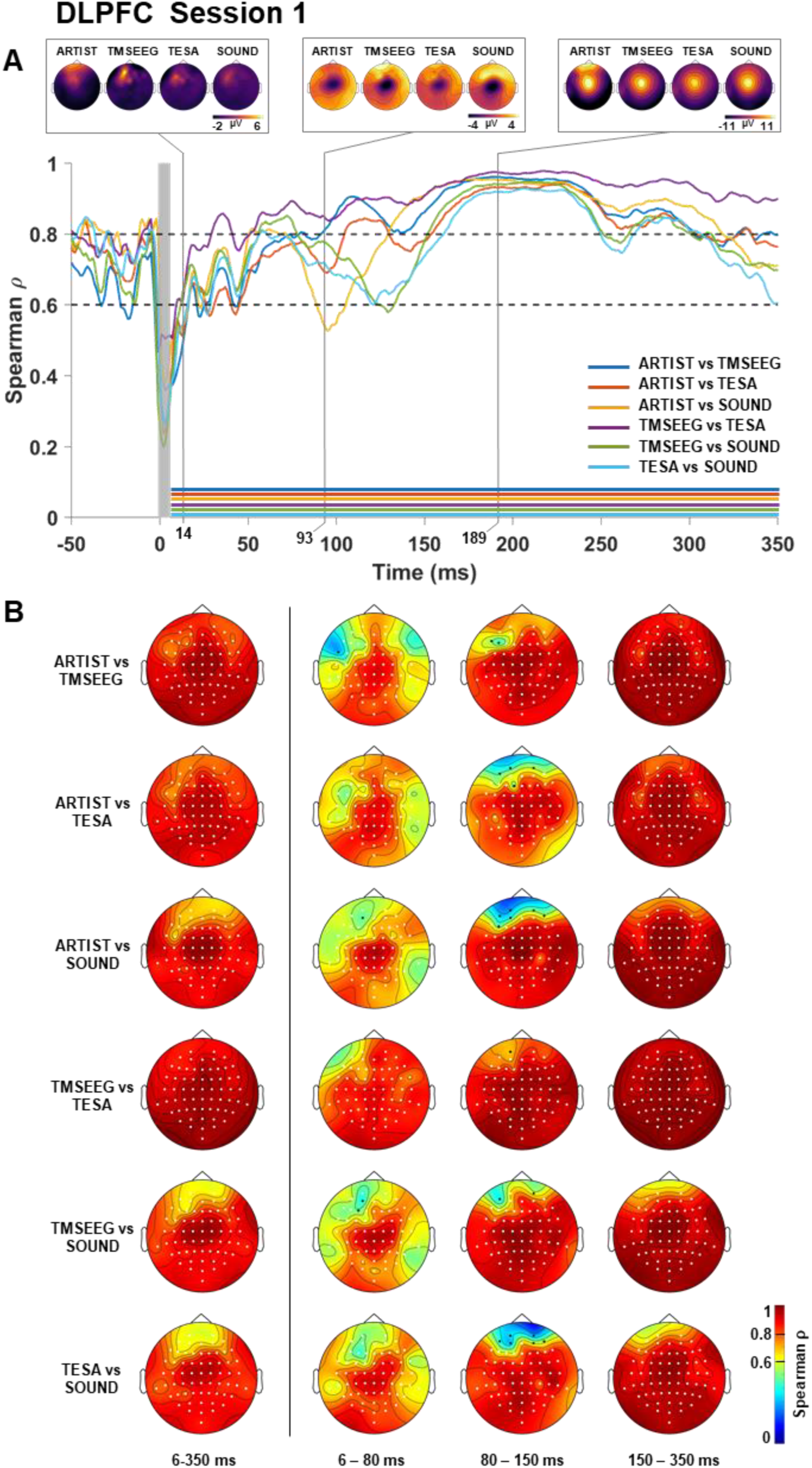
DLPFC, TEPs spatial and temporal correlation. (A) Each colored line represents the spatial correlation (y axis) over time (x axis) of all condition contrasts. Shaded grey column around zero represent the TMS-pulse interpolation interval. Horizontal dotted line represent threshold for moderate (0.6) and substantial (0.8) correlation, according to Shrout et al. 1998. Horizontal colored lines represent instant in time in which the correlation resulted significantly different from zero (p threshold = 0.0042, FDR corrected). On the top, are depicted three representative instantaneous scalp topographies in the four conditions. Voltage on the scalp topographies is color-coded. (B) Temporal correlation of each contrast (rows) in four time intervals (columns). Correlation values are color-coded. Channels significantly different from zero are highlighted in white (p threshold = 0.0042, FDR corrected). “SOUND” refers at the full SOUND-SSP–SIR pipeline.

Accordingly, whole-epoch temporal correlations (Fig. 5-6 B) were moderate-to-substantial (ρ>0.6) for both areas. Correlations at window 6–80 ms were the most variable, showing slight-to-moderate correlation (0.3<ρ<0.6) over frontal and temporal electrodes. TEPs at 80–150 ms showed slight-to-moderate correlations over frontal electrodes in most comparisons (0.2<ρ<0.9), while the rest of the channels were substantially correlated (ρ>0.8). At the 150–350-ms window, TEPs were moderately-to-substantially correlated in all pairs (ρ>0.6).

TEP correlations for session 2 are shown in the supplementary materials (Fig. 5S-6S).

### 3.3 Test–retest reliability comparison across preprocessing pipelines

We identified seven peaks for the IPL (P15, P/N20, P50, N100, P120, P200 and P300) and five peaks for the DLPFC (P20, P50, N100, P200 and N300). Notably, the same peaks were not identifiable in all TEPs: P15 in IPL was found only for SOUND, P120 in IPL only for ARTIST, P50 in IPL was found for ARTIST and TMSEEG while P50 in DLPFC was found only for ARTIST. Figure 7 and Tables 4–5 represent the amplitude and latency CCCs in session 1 vs session 2, for all preprocessing pipelines. CCCs were considered significant when the corresponding bootstrapped CIs did not include zero.

**Table 4:** CCCs (CIs) for amplitude across pipelines in both IPL and DLPFC. Significant values in bold.

| CCC | Amplitude |  |  |  |  |  |  |
| --- | --- | --- | --- | --- | --- | --- | --- |
| IPL |  |  |  |  |  |  |  |
|  | *P15* | *P/N20* | *P50* | *N100* | *P120* | *P200* | *P300* |
| ARTIST |  | 0.46 | 0.37 | -0.10 | **0.64** | **0.73** | **0.55** |
|  |  | (-0.01, 0.77) | (-0.12, 0.72) | (-0.46, 0.28) | **(0.24, 0.86)** | **(0.41, 0.89)** | **(0.10, 0.82)** |
| TMSEEG |  | **0.59** | **0.41** | **0.68** |  | **0.72** | **0.59** |
|  |  | **(0.22, 0.81)** | **(0.03, 0.69)** | **(0.30, 0.87)** |  | **(0.39, 0.89)** | **(0.16, 0.83)** |
| TESA |  | 0.46 |  | **0.54** |  | **0.77** | **0.48** |
|  |  | (-0.02, 0.77) |  | **(0.09, 0.81)** |  | **(0.47, 0.91)** | **(0.05, 0.76)** |
| SOUND | 0.20 | -0.03 | 0.37 | **0.73** |  | **0.69** | 0.47 |
| SSP–SIR | (-0.20, 0.55) | (-0.47, 0.42) | (-0.03, 0.66) | **(0.40, 0.90)** |  | **(0.33, 0.88)** | (-0.01, 0.78) |
| DLPFC |  |  |  |  |  |  |  |
|  | *P20* | *P50* | *N100* | *P200* | *N300* |  |  |
| ARTIST | -0.02 | 0.43 | **0.49** | **0.85** | **0.66** |  |  |
|  | (-0.48, 0.45) | (-0.04, 0.75) | **(0.04, 0.77)** | **(0.63, 0.94)** | **(0.27, 0.87)** |  |  |
| TMSEEG | 0.42 |  | **0.48** | **0.83** | **0.80** |  |  |
|  | (-0.04, 0.73) |  | **(0.09, 0.74)** | **(0.59, 0.93)** | **(0.53, 0.92)** |  |  |
| TESA | 0.14 |  | **0.64** | **0.83** | **0.77** |  |  |
|  | (-0.35, 0.53) |  | **(0.24, 0.86)** | **(0.61, 0.93)** | **(0.50, 0.90)** |  |  |
| SOUND | 0.10 |  | **0.87** | **0.78** | 0.32 |  |  |
| SSP–SIR | (-0.38, 0.53) |  | **(0.70, 0.95)** | **(0.50, 0.91)** | (-0.17, 0.68) |  |  |

**Table 5:**
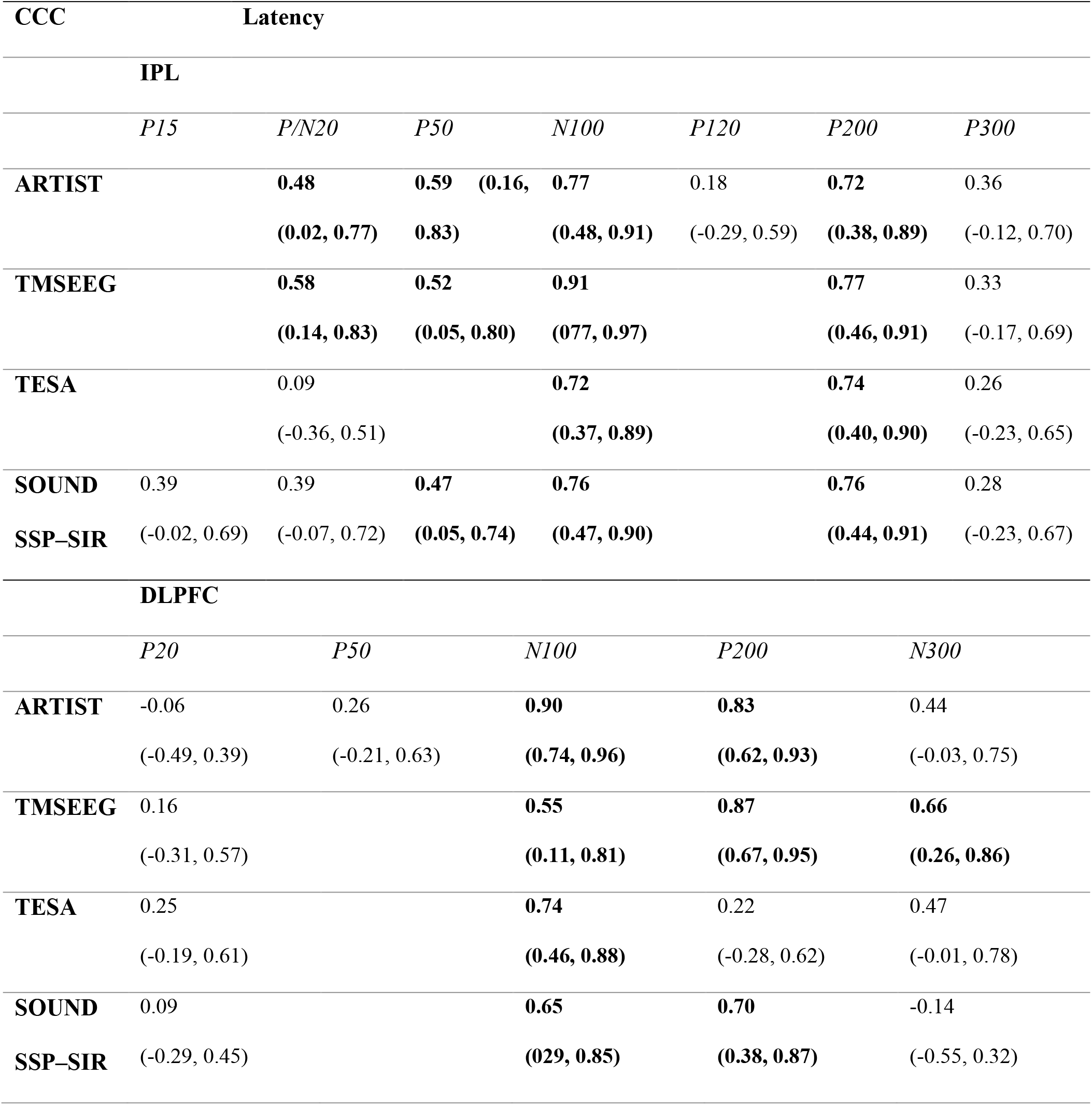
CCCs for latency across pipelines in both IPL and DLPFC. Significant values in bold.

**Fig. 7:**
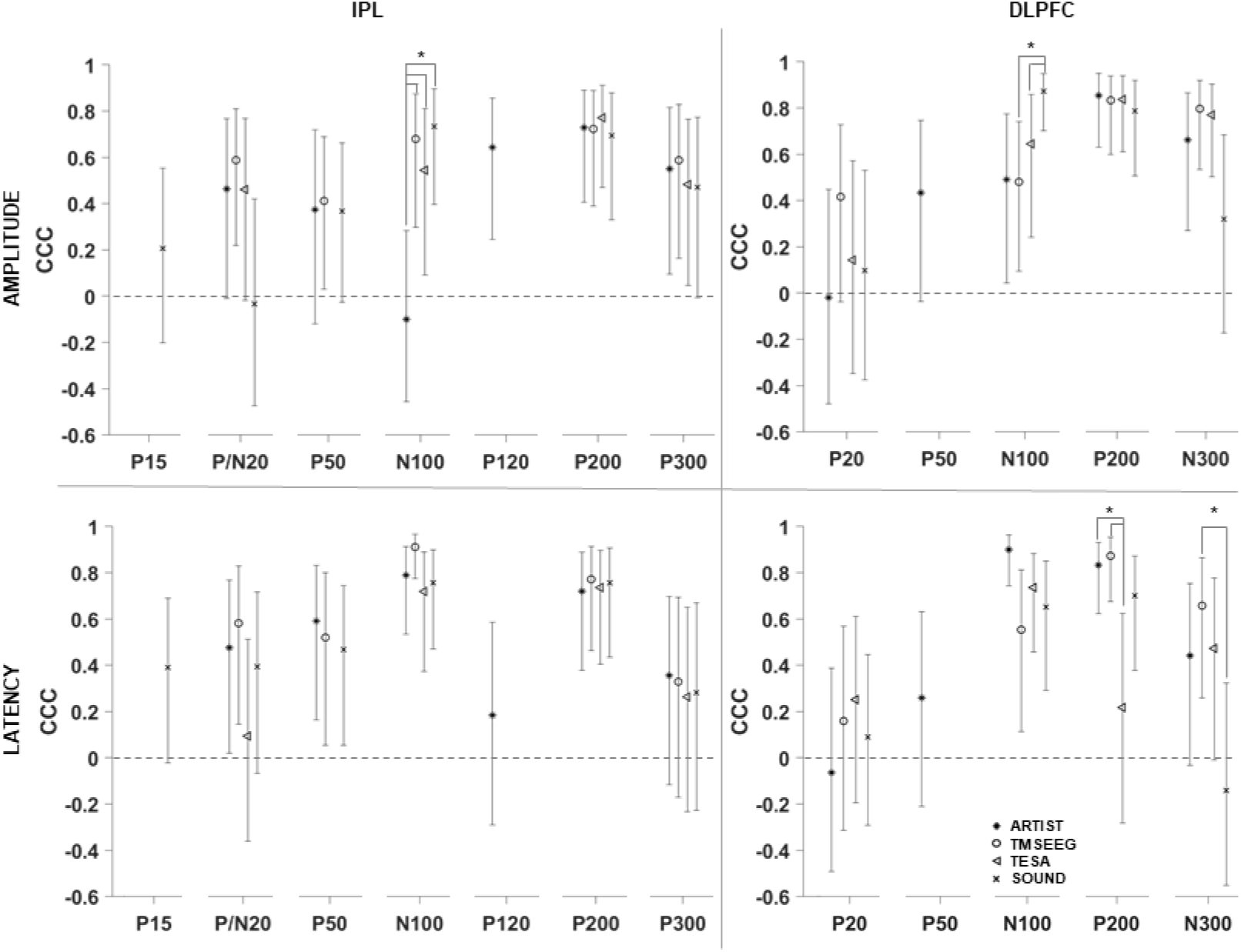
Test-retest reliability of peaks amplitude and latency (rows) in IPL and DLPF (columns), computed on TEPs obtained from the four preprocessing pipelines (see legend). Note that peaks P15, P50 and P120 were not present after all preprocessing pipelines. Vertical error bars represent bootstrapped CIs. CCCs were considered significantly different from zero when the CIs did not include zero (p < 0.05). Asterisks represent significantly different CCCs within the same peak. “SOUND” refers at the full SOUND-SSP–SIR pipeline.

As in the above section, for the interpretation of CCC values we employed the scale in Shrout, 1998 (Shrout, 1998) (Table 5S).

For IPL, CCCs for amplitude were virtually-none-to-fair (−0.03 – 0.59) in early peaks (P15, P/N20, P50) and virtually-none-to-moderate (−0.10 – 0.79) in late peaks (N100, P120, P200 and P300). Notably, the CCC of N100 was significantly different for ARTIST compared to the other preprocessing pipelines (ARTIST vs TMSEEG p=0.002, ARTIST vs TESA *p*=0.004 and ARTIST vs SOUND-SSP–SIR *p*<0.001). Latency CCCs followed a similar pattern (Fig. 7).

For DLPFC, CCC for amplitude was virtually-none-to-fair (−0.02 – 0.43) in early peaks (P20, P50) and increased to slight-to-substantial (0.32 – 0.88) in later peaks (N100, P200 and N300). SOUND-SSP–SIR CCC resulted significantly different in N100 compared to TMSEEG (*p*<0.001) and TESA (*p*=0.022). Latency CCCs in early peaks (P20, P50) were relatively stable around virtually-none-to-slight correlation values (−0.06 – 0.26), while it varied considerably across preprocessing pipelines in later components (N100, P200 and N300), ranging from substantial for ARTIST (0.90) and virtually none for SOUND-SSP–SIR (−0.14). TESA CCC results were significantly different in P200 compared to ARTIST (*p*=0.010) and TMSEEG (*p*=0.036) but not SOUND-SSP– SIR, while TMSEEG CCC resulted significantly different in N300 compared only to SOUND-SSP–SIR (*p*=0.034).

### 3.4 Discussion

We found significant differences in TEP amplitudes across preprocessing pipelines for both stimulated areas, i.e., IPL and DLPFC, in both session 1 and 2. Spatial and temporal correlations were generally moderate-to-substantial, although lower values were found at the beginning of the epoch and on frontal electrodes. Moreover, the test– retest reliability of TEPs was highly variable across components and also varied across preprocessing pipelines. These findings suggest a marked impact of the choice of the preprocessing pipeline on the TEPs, mainly at early latencies, which may affect their reproducibility.

#### Variability brought by the choice of the preprocessing pipeline in the TEPs

The application of different processing pipelines produced TEPs that were significantly different in amplitude, throughout the epoch in IPL and only at late latencies in DLPFC.

Interestingly, TEPs at latencies below and above 100 ms showed different patterns. Above 100 ms, TEPs’ amplitudes were different across pipelines for both areas but also highly correlated with each other in both space and time. This suggests that the functions used to reduce artifacts at those latencies yielded TEPs with similar unfolding of components, but with different amplitudes. Conversely, early-latency TEPs showed less differences in amplitude than late ones, but their correlations were more variable. For example, in the first 15–20 ms after the TMS pulse the spatial correlations were low for most of the comparisons concerning both areas, reaching high values around 95 ms (115 ms for comparisons with ARTIST in the IPL). The temporal correlations in the window 6–80 ms also indicate that most of the low correlations were located in the fronto-temporal regions, where the muscular artifact was more sizeable. It is likely that TEPs before 100 ms are the most affected by the preprocessing, due to the superposition of high-amplitude artifacts such as muscular and decay artifacts. Notice that in our recording the TMS-pulse artifact lasted no longer than 5–6 ms; therefore, it should not have affected the signal after that point.

Late peaks, i.e., N100 and P200, are known as the most reliable and stable peaks of the TEPs (Biabani et al., 2019; Conde et al., 2019; Kerwin et al., 2018; Lioumis et al., 2009; Ozdemir et al., 2020). These studies are in line with our findings in that late latencies were the most resilient to the preprocessing pipeline in terms of shape, resulting in high correlations. However, late peaks are also known to be easily contaminated by the auditory response to the click emitted by the TMS pulse (Biabani et al., 2019; Conde et al., 2019; Nikouline et al., 1999; Nikulin et al., 2003; Tiitinen et al., 1999), unless auditory protection and adequate masking are used to make the click indiscernible. In fact, despite high correlation, we found marked differences in amplitude in late latencies, suggesting a different impact of the preprocessing phase in attenuating those unwanted auditory activations.

These evidences suggest that the preprocessing phase may be a potential source of variability between TEPs, which should be considered when comparing studies that use different preprocessing strategies. For example, beside the N100–P200 complex, TEP components are not yet fully characterized outside the motor cortex. Most of the studies on DLPFC stimulation agree on deflections such as the N40, P60, N100 and P185 (Bagattini et al., 2019; Chung et al., 2019; Conde et al., 2019; Gordon et al., 2018; Rogasch et al., 2014; Voineskos et al., 2019), but other peaks are often reported at ∼30 ms, ∼50 ms, or ∼70 ms (Bagattini et al., 2019; Conde et al., 2019; Rogasch et al., 2014). For IPL, there is still little characterization of TEP components and the few studies that reported them have shown inconsistent results (Conde et al., 2019; Rogasch et al., 2020; Romero Lauro et al., 2014). According to our findings, the preprocessing phase could be a possible source of variability in TEPs’ early-component (<100 ms) definition. It is already known that the peak characterization may be drastically altered should only one TMS-related artifact be left unhandled in the signal (Rogasch et al., 2014). Therefore, the variability in TEP components observed after the data have been preprocessed may be caused by a different artifact suppression in each preprocessing pipeline, especially for large-amplitude early-latency TMS-related artifacts, which are the most difficult to handle.

Moreover, we investigated only four published preprocessing pipelines, while most of the TMS–EEG studies use custom preprocessing pipelines. Therefore, our results could barely represent the unaccounted variability possibly resulting from the various preprocessing pipelines within other TMS–EEG reports.

#### Test–retest reliability estimation of TEPs is affected by the preprocessing pipeline

We found a general trend of low test–retest reliability in early peaks (P15, P20 and P50) and high test–retest reliability in late peaks (N100, P120, P200) (Fig. 6). N/P300 latency was highly variable, probably reflecting the slow response at the end of the epoch. The test–retest reliability of TEPs has been assessed in several studies that are in agreement on the high reliability of late peaks, but reveal also contrasting results for early peaks (Casarotto et al., 2010; Kerwin et al., 2018; Lioumis et al., 2009; Ozdemir et al., 2020). In fact, two studies report high reliability of early peaks (Casarotto et al., 2010; Lioumis et al., 2009) while other two show a reduced reliability in early peaks (Kerwin et al., 2018; Ozdemir et al., 2020). This is in line with our results of spatial and temporal correlations and likely reflect the low SNR in early latencies due to high artifacts such as muscular and decay artifacts associated with fast and small-amplitude cortical components. Conversely, SNR may be higher at late latencies due to higher-amplitude cortical components. Therefore, our findings are in line with the current literature (Kerwin et al., 2018; Ozdemir et al., 2020) that points toward the late components as the most reproducible part of the TEPs and with the greatest potential as a biomarker in clinical research. However, more research is needed to clarify the contribution of the auditory response at those latencies and to extract the direct cortical response to the TMS pulse. Early components may be of interest for the study of effective connectivity as they reflect the activation of remote areas directly connected with the stimulated region. Thus, further methodological studies should be conducted to improve the SNR at these latencies.

Different preprocessing pipelines produced significantly different test–retest reliabilities, in the amplitude of N100 in IPL and DLPFC, and in the latency of P200 and N300 in DLPFC. A major obstacle to this estimation were the large CIs of the CCCs, likely due to our relatively small sample (*N*=16) and high variability of peak amplitudes and latencies across individuals. Nonetheless, we found significant differences between CCCs within the same peak. This result is a practical consequence of the above-mentioned variability associated with the preprocessing phase that adds up to other issues in estimating the test–retest reliability of TEPs. In fact, it is known that TEPs contain sensory responses, i.e., the auditory response, the muscular feedback, the skin receptor feedback, that are time-locked to the pulse and difficult to exclude during the recording or the preprocessing (Biabani et al., 2019; Conde et al., 2019). Several strategies are used to attenuate these contributions, i.e., in-ear headphones playing masking noise, layer of foam applied beneath the coil, to reduce the scalp sensation to the TMS pulse (Farzan et al., 2016). However, these practices do not completely exclude the sensory contribution in TEPs. Therefore, when measuring TEPs’ reliability, to some degree it might not be clear whether we are measuring the reliability of the direct neuronal response or the one from auditory and somatosensory processing, which could itself be a reliable response. This issue could be solved with an appropriate sham control, although this issue continues to be debated in the TMS–EEG field (Belardinelli et al., 2019; Conde et al., 2019; Siebner et al., 2019).

Finally, the dataset itself could have also played a role in generating part of the variability that we described. In fact, TEP signal recorded following DLPFC or IPL stimulation is known to be particularly problematic, because there are not established procedures to set TMS parameters, e.g., TMS intensity and coil orientation, to obtain an optimal cortical response. Moreover, TMS pulses in those areas might interact with head and neck muscles, causing muscular twitches. In our study, the intensity of stimulation was set at 100% MT to minimize the discomfort when stimulating frontal regions, reducing muscular twitches. However, this choice might have contributed to TEPs with low SNR. Therefore, it could be possible that a TMS–EEG dataset recorded in conditions of a higher SNR, like more medial stimulation, with supra-threshold TMS intensities, could have yielded more robust results, less susceptible to the choice of the preprocessing pipeline. Moreover, it should be noted that the interval between the two sessions was relatively long; therefore, our results are relevant in the context of long-interval test–retest reliability, e.g., to develop a neurophysiological biomarker, and may not be relevant for other time intervals. Finally, a few other factors that have not been controlled may have increased the variability of TEPs across sessions (i.e., amount of overnight sleep, follicular cycle, etc.).

#### Possible sources of variability between preprocessing pipelines

The differences observed between the TEP preprocessing pipelines could derive from multiple factors. The main structural differences between the pipelines used in this study were (1) the order of the steps in the pipeline, (2) the type of core functions to remove TMS-related artifacts and (3) the level of automatism.

The order of the steps in processing TMS–EEG signal is crucial, because the interaction of some procedures, i.e., filtering and ICA, with the large-amplitude artifacts might introduce additional analysis-related artifacts to the signal (Rogasch et al., 2017).

Regarding the type of core functions to remove TMS-related artifacts, these methods employ different ICA algorithms [ARTIST uses *Infomax* (Makeig et al., 1997), TMSEEG and TESA use *fastICA* (Hyvärinen and Oja, 2000)] or other methodologies such as the SOUND-SSP–SIR (Mutanen et al., 2016, 2018). *Infomax* and *fastICA* were benchmarked in another study against another ICA algorithm called adaptive mixture ICA (AMICA) (Palmer et al., 2008), which retains the best performance in separating neuronal components so far (Delorme et al., 2012). Both *Infomax* and *fastICA* were highly correlated in terms of component scalp maps and spectral profile with AMICA components. This suggests that using different ICA algorithms in the TEPs should generate minimal variability, since the choice of artifactual components is based on the component scalp maps and spectral profile. However, the selection of ICA, which is done manually by the experimenter in some cases and by automatic algorithms in others, may have impacted the final result. A markedly different approach is the one from SOUND-SSP–SIR. As mentioned in the methods, SOUND cleans the signal in each sensor at the source level utilizing the information coming from nearby sensors, while SSP–SIR attenuates the TMS-induced muscular artifact detecting its time–frequency features at the source level. Another study already compared ICA and SSP–SIR (without SOUND) approaches in recovering the TMS–EEG neuronal signal (Biabani et al., 2019). There, ICA was found to reduce and distort the signal more than SSP–SIR. In contrast with our expectation, we found that SOUND-SSP– SIR’ TEPs did not show marked statistical differences with all the ICA-based methodologies. Nonetheless, ICA might not be the ideal tool to separate brain signal from TMS-related artifact, because one of its core assumptions is that these two signals are independent from each other. Although this assumption might hold in the EEG alone, in TMS–EEG many artifacts are time-locked to the TMS pulse, as it is the brain signal, breaking the assumption of independency (Metsomaa et al., 2014). This might be an issue for the ICA-based preprocessing pipelines.

As mentioned in section 2, the level of automation was substantially different across pipelines. ARTIST was designed to be fully automatic, in order to minimize the error due to subjective choices of the experimenter. Therefore, trials, channels and ICA components were rejected without supervision. In our study, this feature led to a substantial difference in the number of channels and ICA components rejected from this methodology. This could explain why ARTIST TEPs resulted as the most different in terms of amplitude in both IPL and DLPFC and also why spatial correlations with ARTIST deviated from the others until almost 125 ms in the IPL. TMSEEG, TESA and SOUND-SSP–SIR also showed significant differences in rank after the preprocessing between each other, although remarkably less pronounced than ARTIST. This is also reflected in the higher spatial and temporal correlations and in the reduced differences of the cluster-based ANOVA between those pipelines. Trial removals were different between pipelines. Specifically, SOUND-SSP–SIR pipeline consistently removed less trials than the other pipelines. This is not surprising, since SOUND-SSP–SIR approach is to clean the data before doing the trial rejection, in order to spare the signal as much as possible.

#### What to do to overcome the variability brought by the preprocessing approach?

Variability brought by differences in the preprocessing phase is a crucial emerging issue that we have reported here for TMS–EEG and that has been recently reported also for other techniques applied in the neuroscience field, such as fMRI (Botvinik-Nezer et al., 2020; Lindquist, 2020). Importantly, this variability on TMS–EEG studies would be minimized when the results are based on the contrast between two conditions, processed with the same pipeline, e.g., TEPs pre vs post treatment or TEPs rest vs task, because the preprocessing phase would likely affect the signal in the same way in both conditions. However, in studies on resting-state TMS–EEG aimed to investigate effective connectivity, the choice of the preprocessing phase would be more critical, since there would be no contrast to control its effect. The main obstacle to solve this issue is the lack of a ground truth, i.e., a clean signal, that would allow one to assess the efficacy of each preprocessing pipeline. One way could be to process simulated TMS–EEG data, where the ground truth is known. Although there are technical difficulties in reconstructing a plausible TMS–EEG signals, with the TMS-related artifacts, there has been recent development in simulating physiologically realistic EEG signals (Neymotin et al., 2020), which could be potentially combined with, e.g., realistic-sham-elicited brain and scalp-muscle responses. Furthermore, some methods, such as linear spatial filtering methods like SOUND and SSP–SIR, allow a straightforward quantification of the over correction across all the possible cortical sources (Mutanen et al., 2016). Because the spatial filters are perfectly known, one can apply them to all lead-field topographies to determine which sources would be attenuated the most by, e.g., SOUND or SSP–SIR. This approach enables the assessment of potential distortions in the effective connectivity patterns of interest and the suitability of the chosen analysis pipeline for the research question.

In the absence of a proper benchmark, good practices would be similar to the ones proposed by Botvinik-Nezer et al. (2020) for fMRI: the TMS–EEG field would surely benefit from the sharing of code and data, ultimately allowing for a more straightforward replication of the findings. The use of pre-registration, in which the selection of parameters are explicated, is another practice that would reduce the risk of cherry-picking the preprocessing steps that best suit the original hypothesis. Moreover, when there are no *a priori* hypotheses on a particular preprocessing pipeline, the use of a multiverse analysis (Steegen et al., 2016), i.e., the use of multiple statistical and analytical approaches in the same study to reach consensus findings, would be a way to control the variability intrinsic to the preprocessing phase, yielding more solid results.

## Acknowledgments

This work was supported by the Center of Mind/Brain Sciences - CIMeC – University of Trento by Fondazione Caritro, by the Italian Ministry of Health – Ricerca Corrente, and the Academy of Finland (Grant No. 321631).

