## Supplementaty material for "The impact of artifact removal approaches on TMS–EEG signal"

### Supplementary methods

The study from which the sample comes from Esposito et al. (in preparation) included three sessions, one for MRI acquisition and two TMS-EEG recordings. Structural and functional imaging at rest were recorded for each subject. From these data, cortical nodes of the DMN, which were used as target areas for the TMS-EEG sessions, were extracted individually for each subject. The left pre-frontal and parietal nodes were identified through independent component analysis (ICA) of the individual resting-state fMRI. Firstly, a group ICA using MELODIC within FSL 5.0.9 [1] was performed to extract the group DMN. The target network was chosen based on a template. Pre-processed resting-state fMRI data were normalized to the MNI template and decomposed into ten independent components [2] and then thresholded at z-scores > 2.3, $P<0.05$. Subsequently, a dual-regression was used to derive single subject DMN from the group DMN [3,4]. Single-subject DMN volume maps were thresholded at $z > 2.3$, $P < 0.05$. For each subject, the left prefrontal and parietal cortical nodes of the DMN were manually extracted and used as TMS targets.

### Supplementary results

Fig. 1S shows TEPs with relative topographies after being processed with the four methods, in both IPL (Fig. 1S A) and DLPFC (Fig. 1S B) for session 2.

The ANOVAs on average percentage of removed trials (table 1S) in session 2 were significant for both IPL ($\chi^2$ = 13.1850, p = 0.0046) and DLPFC ($\chi^2$ = 10.8589, p = 0.0125).The post-hoc tests did not reveal any significant difference.

The ANOVAs on data rank after the preprocessing in session 2 were significant for the IPL (F= 113.651, p < 0.001) and DLPFC session 2 (F= 130.475, p < 0.001). Post-hoc results are illustrated in table 2S and figure 2S.

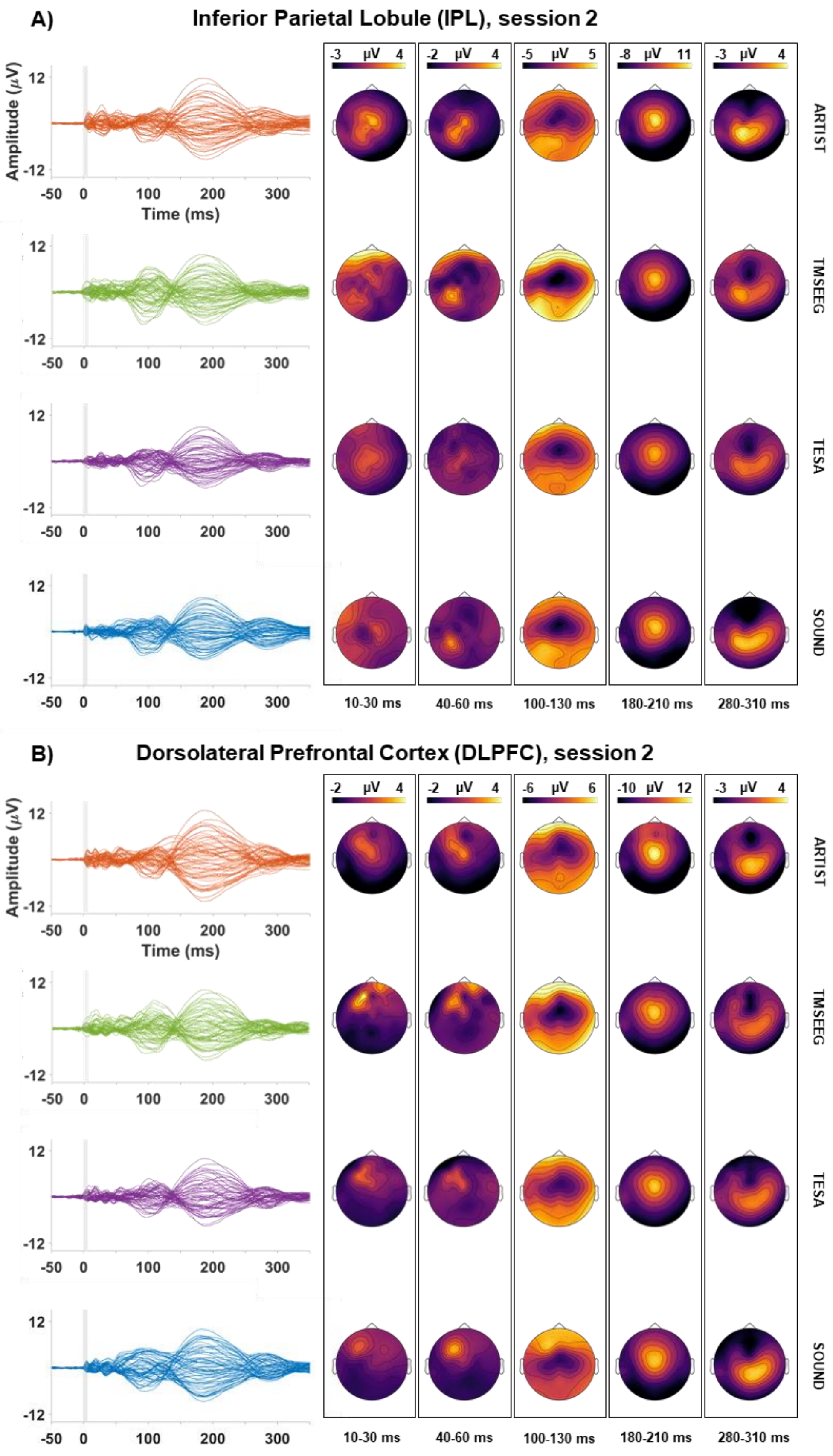


Fig.1S: TEPs recorded after IPL (A) and DLPFC (B), processed with different pipelines (rows). The leftmost column depicts the TEPs time-series, color-coded for each preprocessing pipeline. The other columns represent the scalp topographies of the TEPs in five selected intervals. The color on the topographies represent the mean voltage on the scalp in each interval. *"SOUND" refers at the full SOUND-SSP–SIR pipeline.*

*Table 1S: trials removed by each preprocessing method (upper row: mean (SD), lower row: median (range))*

| | **Session 1** | | **Session 2** | |
|---|---|---|---|---|
| | *IPL* | *DLPFC* | *IPL* | *DLPFC* |
| **ARTIST** | 7.94 (4.96) | 6.75 (4.41) | 6.13 (3.81) | 8.38 (5.23) |
| | 7.5 (16-0) | 4.5 (17-2) | 6 (19-0) | 9 (20-0) |
| **TMSEEG** | 8.38 (8.25) | 11.13 (6.85) | 4.31 (4.44) | 5.75 (4.88) |
| | 5 (34-0) | 12 (24-1) | 3 (10-2) | 3.5 (18-0) |
| **TESA** | 7.50 (4.36) | 6.56 (3.06) | 5.50 (2.26) | 6.25 (2.49) |
| | 6 (18-1) | 6 (13-2) | 6 (10-2) | 5 (13-4) |
| **SOUND** | 2.50 (2.87) | 3.44 (2.71) | 2.86 (3.47) | 3.44 (3.00) |
| **SSP-SIR** | 1 (8-0) | 4 (8-0) | 1 (13-0) | 3 (10-0) |

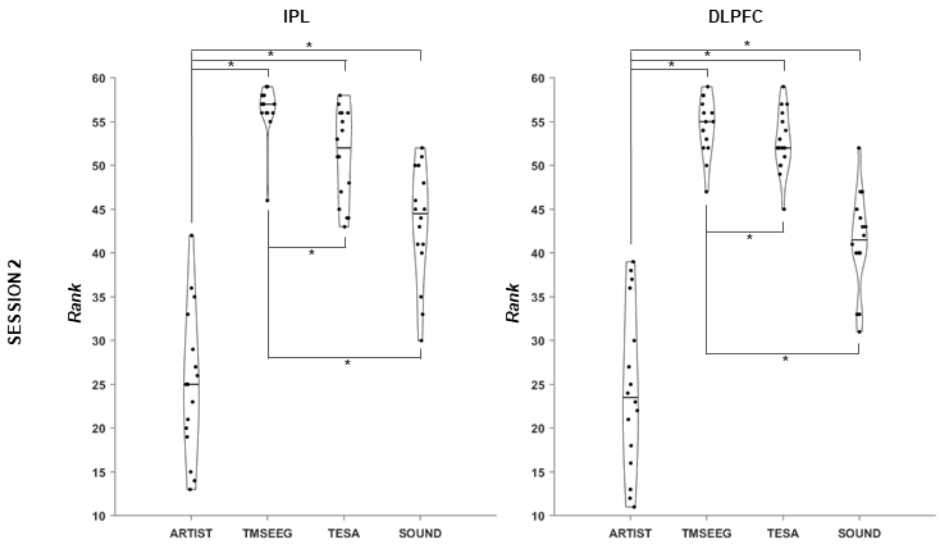


*Fig. 2S: Data rank after each preprocessing methods in the four conditions. Each violin plot depicts the distribution of individual subject rank after the preprocessing. Horizontal black lines represent the median. Black lines with starts connects conditions significantly different in the post-hoc analysis (p threshold= 0.0042). "SOUND" refers at the full SOUND-SSP–SIR pipeline.*

*Table 2S: Post-hoc tests on data rank across preprocessing methods in each condition. Significant differences in bold.*

| | | IPL session 1 | | DLFPC session 1 | | IPL session 2 | | DLPFC session 2 | |
|---|---|---|---|---|---|---|---|---|---|
| **ARTIST** | TMSEEG | **t = -18.711** | **p < 0.001** | **t = -21.205** | **p < 0.001** | **t = -17.256** | **p < 0.001** | **t = -17.576** | **p < 0.001** |
| | TESA | **t = -15.913** | **p < 0.001** | **t = -19.189** | **p < 0.001** | **t = -14.294** | **p < 0.001** | **t = -16.482** | **p < 0.001** |
| | SOUND | **t = -11.157** | **p < 0.001** | **t = -13.066** | **p < 0.001** | **t = -10.023** | **p < 0.001** | **t = -9.809** | **p < 0.001** |
| **TMSEEG** | TESA | t = 2.798 | p = 0.045 | t = 2.016 | p = 0.299 | t = 2.962 | p = 0.029 | t = 1.094 | p = 1.000 |
| | SOUND | **t = 7.554** | **p < 0.001** | **t = 8.138** | **p < 0.001** | **t = 7.233** | **p < 0.001** | **t = 7.767** | **p < 0.001** |
| **TESA** | SOUND | **t = 4.756** | **p < 0.001** | **t = 6.123** | **p < 0.001** | **t = 4.271** | **p = 0.001** | **t = 6.673** | **p < 0.001** |

### Differences in TEPs amplitude (session 2)

For IPL, TEPs derived from the four preprocessing methods were significantly different in amplitude (IPL: p =0.001, cluster window = 6 - 350 ms), as shown by the cluster-based ANOVA. In post-hoc tests (Fig. 3S), significant clusters were found in five contrast over six. Only the contrast for TMSEEG vs TESA did not show significant differences (Table 3S).

Similarly, TEPs for DLPFC stimulation were different across processing pipelines (DLPFC: p =0.001, cluster window = 10 – 350 ms), as shown by the cluster-based ANOVA. In post-hoc tests (Fig. 4S), significant clusters were found in all contrasts.

Table 3S: IPL post-hoc paired t-tests, session 2

| Comparison | Cluster Polarity | Cluster p | Cluster Latency (s) |
|---|---|---|---|
| **ARTIST vs TMSEEG** | + | < 0.001 | 0.014 to 0.350 |
| **ARTIST vs TESA** | + | < 0.001 | 0.151 to 0.350 |
| **ARTIST vs SOUND SSP-SIR** | + | < 0.001 | 0.117 to 0.350 |
| | - | < 0.001 | 0.189 to 0.350 |
| **TMSEEG vs TESA** | | / | |
| **TMSEEG vs SOUND SSP-SIR** | - | < 0.001 | 0.197 to 0.350 |
| **TESA vs SOUND SSP-SIR** | - | 0.002 | 0.199 to 0.350 |

Table 4S: DLPFC post-hoc paired t-tests

| Comparison | Cluster Polarity | Cluster p | Cluster Latency (s) |
|---|---|---|---|
| **ARTIST vs TMSEEG** | + | < 0.001 | 0.012 to 0.350 |
| **ARTIST vs TESA** | + | < 0.001 | 0.155 to 0.350 |
| **ARTIST vs SOUND SSP-SIR** | + | < 0.001 | 0.183 to 0.350 |
| | - | < 0.001 | 0.187 to 0.350 |
| **TMSEEG vs TESA** | + | 0.0030 | 0.080 to 0.350 |
| **TMSEEG vs SOUND SSP-SIR** | - | 0.0030 | 0.208 to 0.350 |
| **TESA vs SOUND SSP-SIR** | - | 0.0040 | 0.218 to 0.350 |

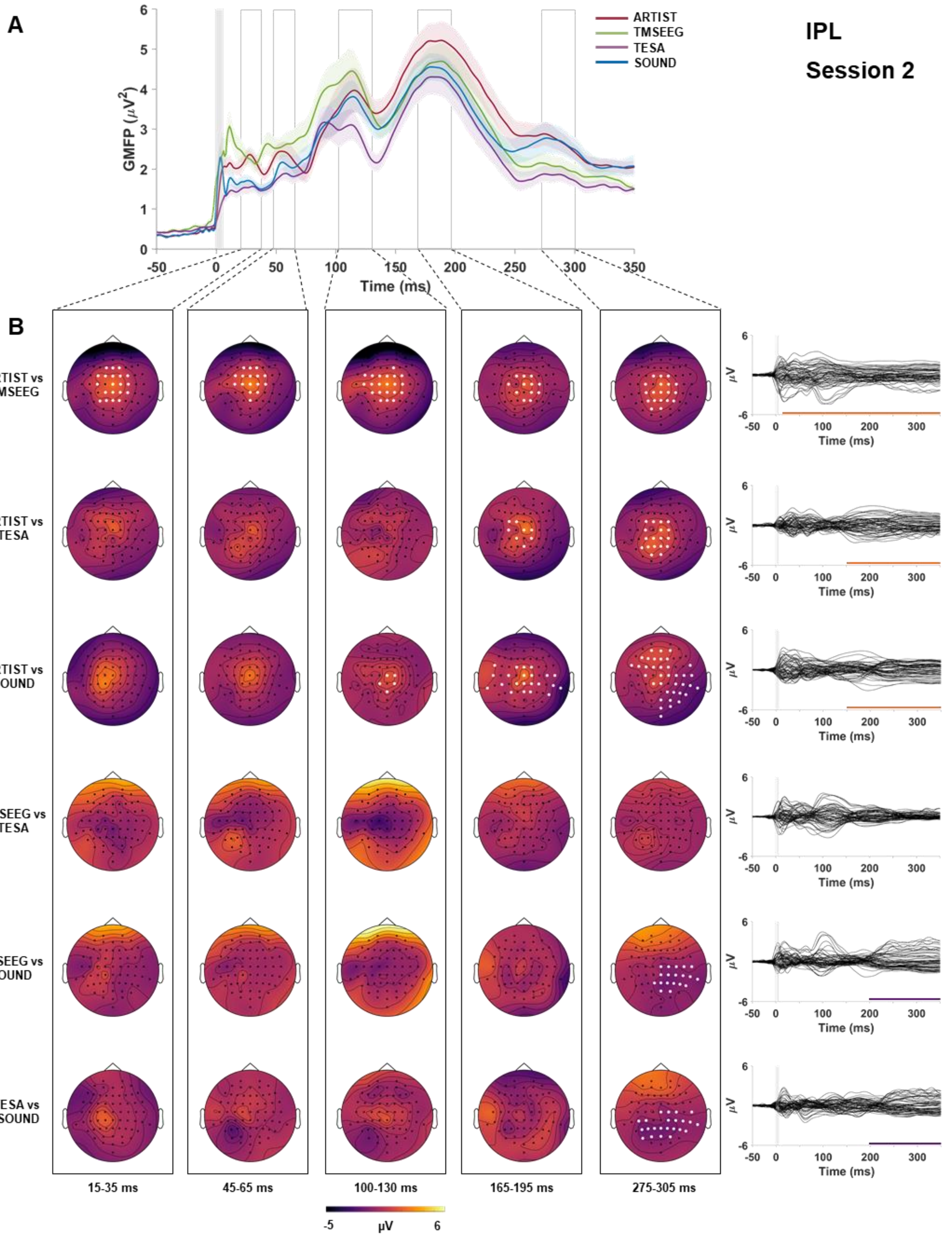

A
IPL
Session 2
ARTIST
TMSEEG
TESA
SOUND
GMFP (μV²)
Time (ms)
B
ARTIST vs TMSEEG
ARTIST vs TESA
ARTIST vs SOUND
TMSEEG vs TESA
TMSEEG vs SOUND
TESA vs SOUND
15-35 ms
45-65 ms
100-130 ms
165-195 ms
275-305 ms
μV
-5
6

*Fig.5S: A: Global Mean Field Power (GMFP, y axes) over time (ms, x axis) of the TEPs resulting from IPL stimulation, cleaned with the four preprocessing pipelines (color-coded). Shaded area around each colored line represents SEM. Shaded grey column around zero represents the TMS-pulse interpolation interval. B: scalp topographies of the voltage differences (color-coded) for each condition contrast (rows) in five selected time windows (columns). White dots represent significant channels. The rightmost column represent the time-series of the voltage differences over time in each contrast. Colored bars at the bottom of each time-series represent the temporal extend of significant cluster(s). Orange positive clusters, purple negative cluster. "SOUND" refers at the full SOUND-SSP–SIR pipeline.*

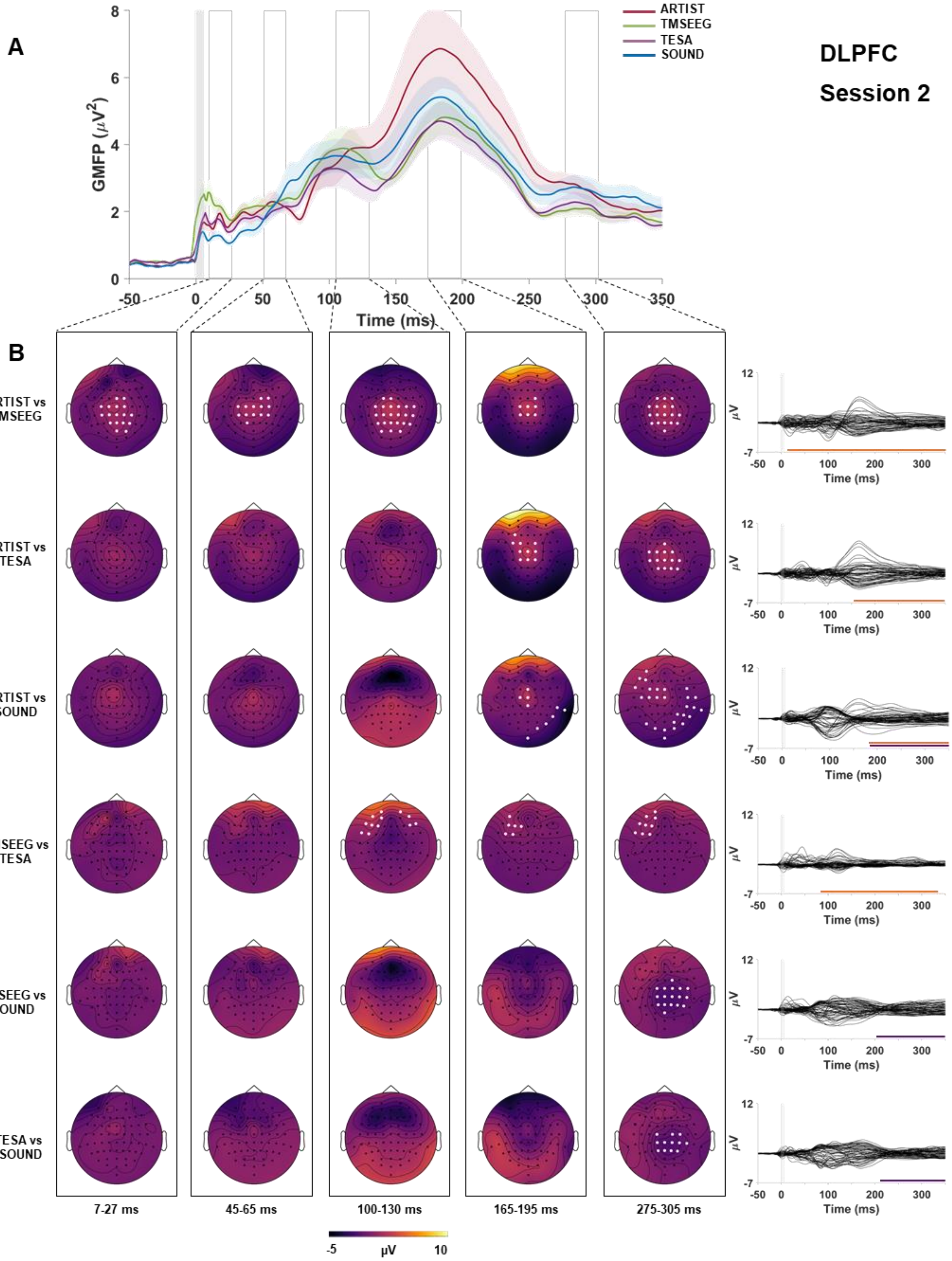
A
DLPFC
Session 2
ARTIST
TMSEEG
TESA
SOUND
GMFP (μV²)
Time (ms)
B
ARTIST vs TMSEEG
ARTIST vs TESA
ARTIST vs SOUND
TMSEEG vs TESA
TMSEEG vs SOUND
TESA vs SOUND
7-27 ms
45-65 ms
100-130 ms
165-195 ms
275-305 ms
-5
μV
10

*Fig.4S: A: Global Mean Field Power (GMFP, y axes) over time (ms, x axis) of the TEPs resulting from IPL stimulation, cleaned with the four preprocessing pipelines (color-coded). Shaded area around each colored line represents SEM. Shaded grey column around zero represents the TMS-pulse interpolation interval. B: scalp topographies of the voltage differences (color-coded) for each condition contrast (rows) in five selected time windows (columns). White dots represent significant channels (p threshold = 0.0042, cluster based corrected). The rightmost column represent the time-series of the voltage differences over time in each contrast. Colored bars at the bottom of each time-series represent the temporal extend of significant cluster(s). Orange positive clusters, purple negative cluster. "SOUND" refers at the full SOUND-SSP–SIR pipeline.*

Table 5S: Landis and Koch adjectives, revisited by Shrout et al. 1998 [5]

| Correlation value | Interpretation |
|---|---|
| **0.00, 0.10** | Virtually none |
| **0.11, 0.40** | Slight |
| **0.41, 0.60** | Fair |
| **0.61, 0.80** | Moderate |
| **0.80, 1,00** | Substantial |

**TEPs correlations (session 2)**

For interpretation of correlation values we employed the scale in Shrout, 1998 [5] (Table 5S).

As in session 1, spatial correlations in the baseline period were high for both areas (moderate-to-substantial, ρ 0.6-0.9). Spatial correlation dropped around the TMS pulse, and then slowly recovered. Correlation values at early latencies (< 100 ms) were the most variable across comparisons ranging from approximately ρ 0.09 to 0.9, while correlations at later (> 100 ms) latencies were moderate-to-substantial (ρ>0.6), consistently in all pairs (fig. 5S-6S A).

Whole-epoch temporal correlations (fig. 5S-6S B) were moderate-to-substantial (ρ>0.6) for both areas, with lowest values being located on frontal electrodes. Correlations at 6-80 ms and 80-150 ms showed slight-to-moderate correlation (0.3<ρ<0.6) over frontal and temporal electrodes and moderate-to-substantial correlation (ρ < 0.6) over the rest of the scalp. TEPs at late-latencies window (150-350 ms) were substantially correlated in most pairs (ρ>0.8).

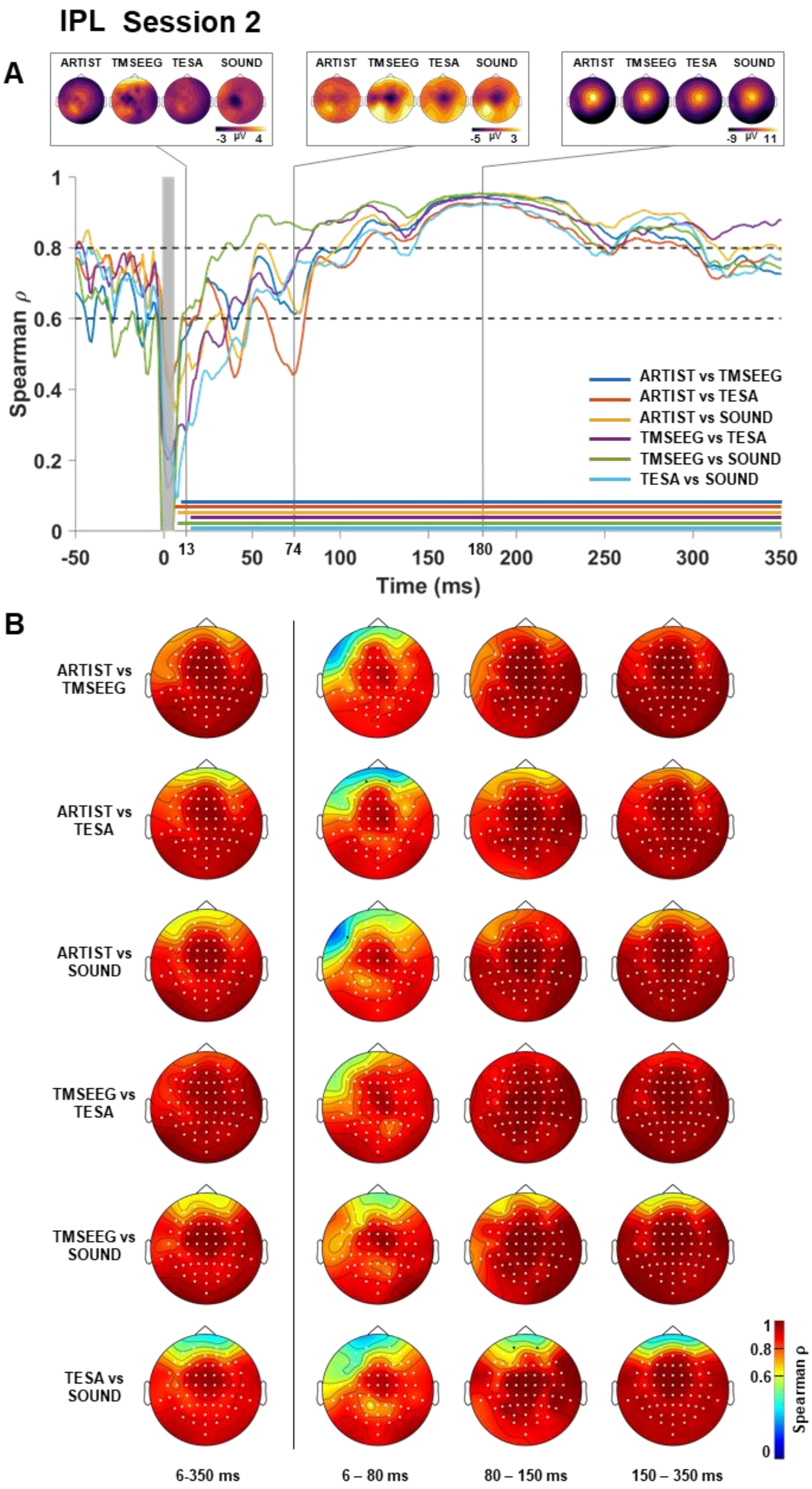


Fig.5S: IPL, TEPs spatial and temporal correlation. (A) Each colored line represent the spatial correlation (y axis) over time (x axis) of all condition contrasts. Shaded grey column around zero represent the TMS-pulse interpolation interval. Horizontal dotted line represent threshold for moderate (0.6) and substantial (0.8) correlation, according to Shrout et al. 1998. Horizontal colored lines represents instant in time in which the correlation resulted significantly different from zero (p threshold = 0.0042, FDR corrected). On the top, are depicted three representative instantaneous scalp topographies in the four conditions. Voltage on the scalp topographies is color-coded. (B) Temporal correlation of each contrast (rows) in four time intervals (columns). Correlation values are color-coded. Channels significantly different from zero are highlighted in white (p threshold = 0.0042, FDR corrected). "SOUND" refers at the full SOUND-SSP–SIR pipeline.

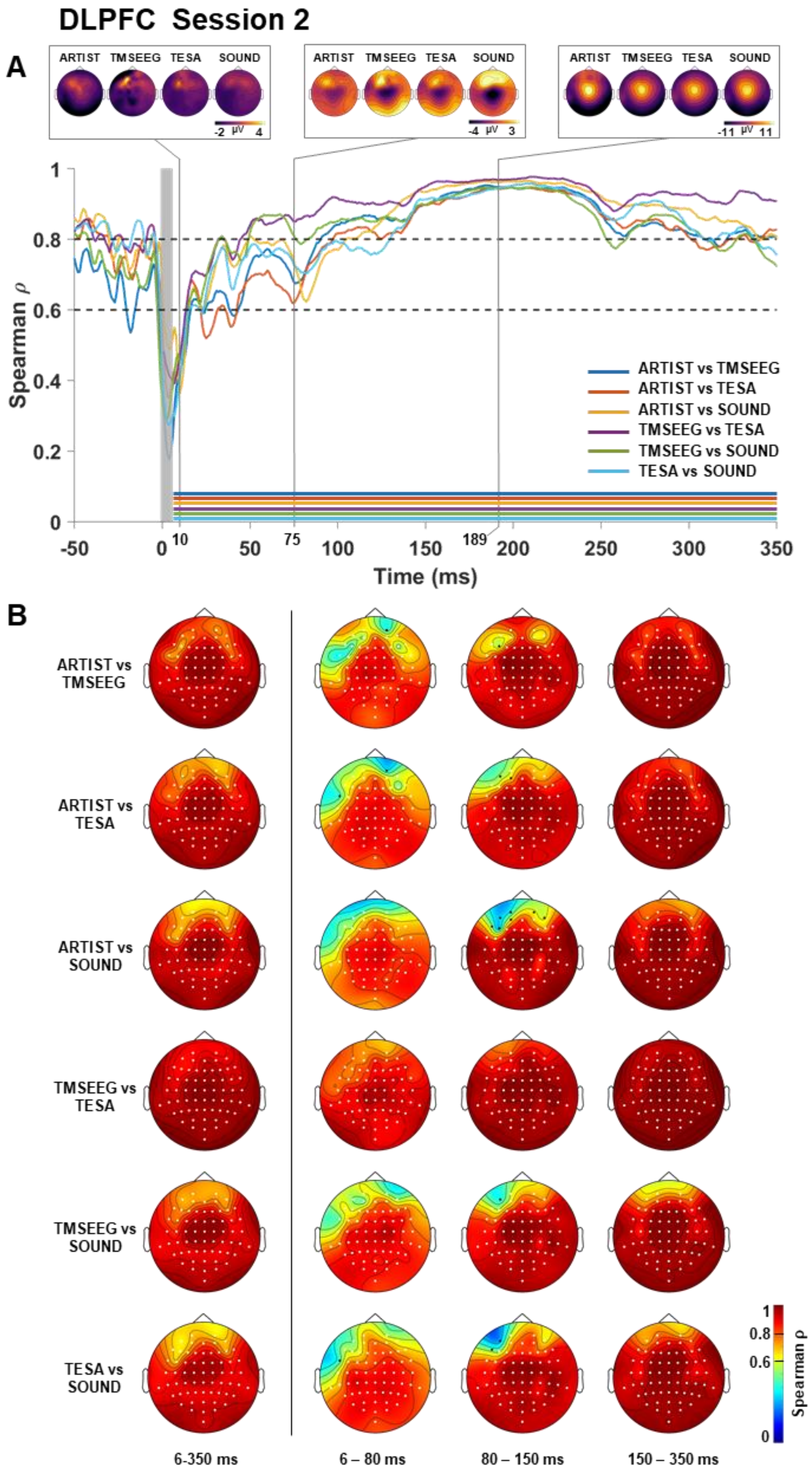


*Fig.6S: DLPFC, TEPs spatial and temporal correlation. (A) Each colored line represent the spatial correlation (y axis) over time (x axis) of all condition contrasts. Shaded grey column around zero represent the TMS-pulse interpolation interval. Horizontal dotted line represent threshold for moderate (0.6) and substantial (0.8) correlation, according to Shrout et al. 1998. Horizontal colored lines represents instant in time in which the correlation resulted significantly different from zero (p threshold = 0.0042, FDR corrected). On the top, are depicted three representative instantaneous scalp topographies in the four conditions. Voltage on the scalp topographies is color-coded. (B) Temporal correlation of each contrast (rows) in four time intervals (columns). Correlation values are color-coded. Channels significantly different from zero are highlighted in white (p threshold = 0.0042, FDR corrected). "SOUND" refers at the full SOUND-SSP–SIR pipeline.*
